# The membrane transition strongly enhances biopolymer condensation through prewetting

**DOI:** 10.1101/2024.08.26.609758

**Authors:** Yousef Bagheri, Mason Rouches, Benjamin Machta, Sarah L. Veatch

## Abstract

Biopolymers that separate into condensed and dilute phases in solution also prewet membranes when one or more components couple to membrane lipids. Here we demonstrate that this prewetting transition becomes exquisitely sensitive to lipid composition when membranes have compositions near the boundary of liquid-ordered/liquid-disordered phase coexistence. In simulation and reconstitution, we couple polyelectrolytes to membranes of saturated lipids, unsaturated lipids, and cholesterol, and find that the coexistence of prewet and dry surface phases is dramatically potentiated by proximity to the membrane phase transition. In cells, we employ an optogenetic tool to characterize prewetting both at the plasma membranes (PM) and the endoplasmic reticulum (ER), and find that prewetting is potentiated or inhibited by perturbations of membrane composition. Prewetting can also mediate membrane adhesion, with avidity dependent on membrane composition. This effect is demonstrated in cells through the potentiation or inhibition of ER-PM contact sites. The strong correspondence between results in simulation, reconstitution, and cells demonstrates a new role for membrane lipids in regulating the recruitment and assembly of soluble proteins.

## INTRODUCTION

Membranes often act as assembly sites for soluble proteins. These structures can be relatively stable, scaffolding the protein machinery needed to perform biological functions, for example at focal adhesions (1), neurological synapses (2), or at tight junctions (3). These structures can also be assembled in response to dynamic stimuli, gathering proteins that propagate signals, for example the signaling platforms that form downstream of T cell receptor activation (4). In a growing number of cases, protein assembly at membranes is attributed to phase separation, driven by multivalent interactions between specific proteins (5–14). This occurs when protein combinations capable of condensing in solution are localized to membranes, often by tethering one or more components to the membrane or by including proteins that interact with lipids. Recent work has shown that localizing condensates to membranes can drive a range of phenomena in reconstituted systems, including membrane curvature, lipid ordering, shifting phase transition temperatures, and transbilayer communication (15–21). Often overlooked is the possibility that membranes themselves can contribute to protein condensation, by tuning interactions between tethering components.

We recently developed a theory of biopolymer prewetting at membranes (22). Unlike wetting, which describes how preexisting phase separated domains spread and deform at surfaces, prewetting describes condensation at surfaces when coexistence is thermodynamically unstable in the bulk. Classical prewetting describes the condensation of liquids at solid surfaces (23, 24), and is observed within a narrow region close to the gas-liquid transition (25). Recent theory predicts that the prewetting region expands when the surface is fluid (22, 26), in good agreement with experimental observations of protein and/or nucleic acid condensation at membranes (5–9). Our theory additionally predicts that the prewetting regime can be further expanded in the presence of membrane-mediated interactions between tethers, for example those that arise close to a membrane phase transition (22). Since mammalian cell plasma membranes have compositions tuned to near phase separation (27, 28), prewetting could provide a sensitive means to organize proteins at the cell surface.

The goal of the current study is to experimentally probe the role of the membrane phase transition in tuning prewetting at membranes in controlled reconstituted and cellular systems. This work systematically explores the regions of phase space where prewetting is most impacted, sometimes dramatically, by interactions within the membrane plane, extending upon recent studies that document coupling between membrane and protein phase transitions in select contexts (11, 21). Under these conditions, perturbations that exclusively target membrane components can control the recruitment and condensation of soluble factors and the avidity of adhesion between membranes, suggesting new roles for membrane properties in broad cellular processes.

## RESULTS

### Prewetting of polyelectrolytes on a fluid lipid bilayer

We first explore a reconstituted system, utilizing a simplified model consisting of short poly-L-lysine (pLys) and poly-L-glutamic acid (pGlu) polyelectrolytes coupled to a supported planar membrane. pGlu and pLys mixtures are a well characterized system that undergoes complex coacervation (29). Figure 1A is a schematic representation of our experimental approach. We mix soluble pLys and pGlu polyelectrolytes in solution at an equal weight ratio in the presence of phosphate buffer (pH=7.4) and NaCl to tune the magnitude of electrostatic interactions. In the absence of a membrane, these mixtures separate into polymer condensed and dilute liquid phases over a range of polymer and NaCl concentrations (29). We also include a fluid membrane, in this case made up of a single unsaturated lipid (DOPC) along with a small fraction (4 mole%) of a reactive DOPE-DBCO lipid to facilitate covalent conjugation of an azide conjugated pLys (N_3_-pLys) through click chemistry (30). pLys-conjugated lipid tethers (DOPE-pLys) couple soluble polyelectrolytes to the membrane. Bilayer membranes are glass supported multilayers assembled by hydrating spin-coated lipid films to reduce the impact of the substrate (31).

**Figure 1:**
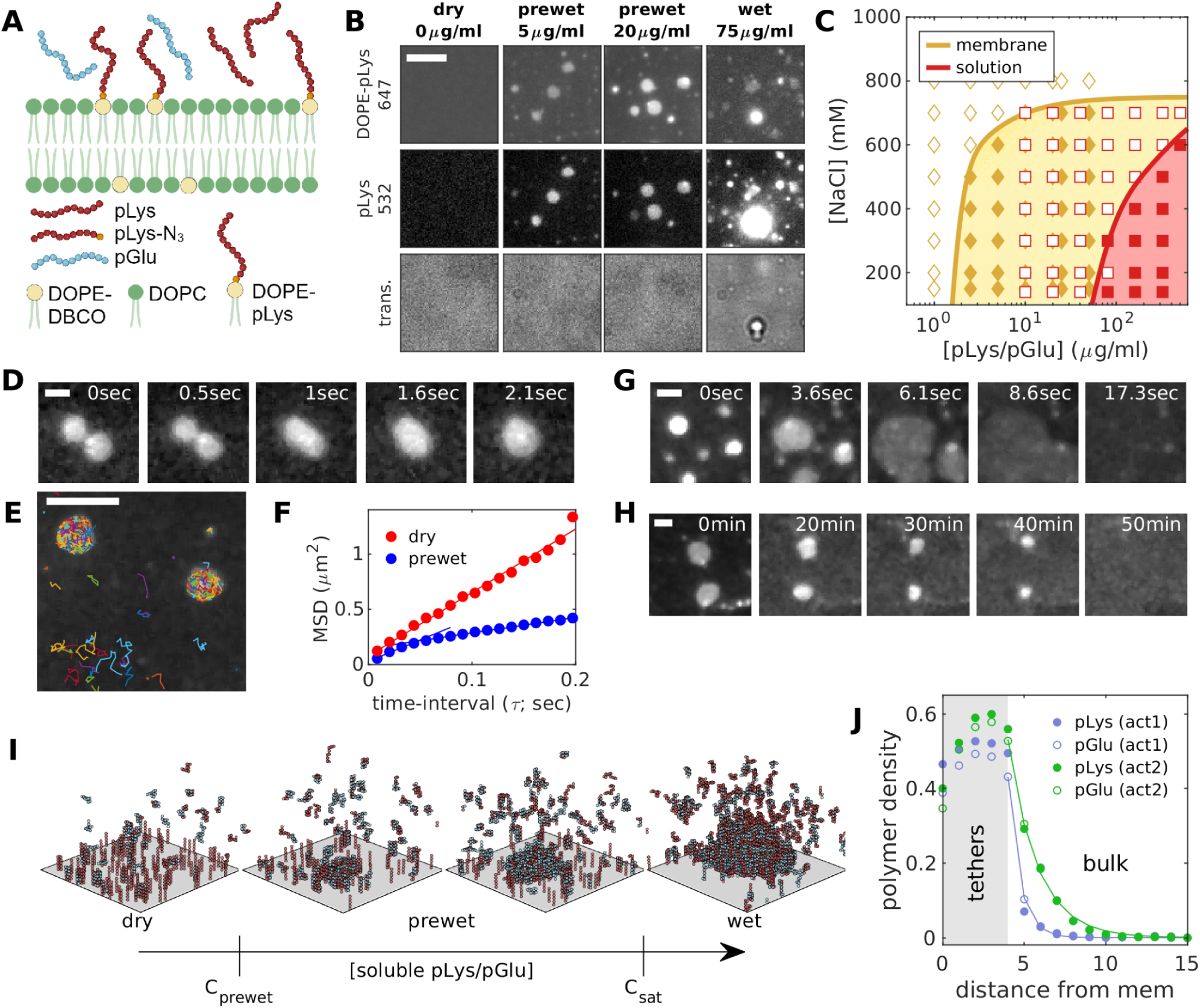
polyelectrolyte prewetting at simple membranes. (A) Schematic representation of the experimental system containing bulk pLys and pGlu polyelectrolytes in contact with a DOPC membrane with DOPE-pLys tethers. (B) Membranes exposed to varying concentrations of 1:1 w/w pLys/pGlu polyelectrolytes in the presence of 400mM NaCl. Prewet domains containing tethers (DOPC-pLys-647) and soluble polymer (pLys-532) are observed at intermediate polyelectrolyte concentrations (5 and 20μg/ml) while wetting is detected at high concentrations (75μg/ml). Wet but not prewet domains visible in transmitted light images. (C) Phase diagram marking polyelectrolyte and NaCl concentration where prewet (yellow) or bulk (red) domains are detected in microscopy images. (D) Prewet domains are dynamic, circular, and can coarsen via coalescence. Images are snapshots from Supplementary Movie 1. (E) Single molecule trajectories of tethers reveal they are dynamic and (F) plots of the mean squared displacement (MSD) vs time-interval for domain localized tethers are sub-linear, indicating they are confined. (G) Prewet polyelectrolyte domains dissipate when exposed to high NaCl concentration (2M) and when (H) soluble polyelectrolytes are removed from solution. Images are snapshots from Supplementary Movies 2,3. Scale bar is 5μm in B,E and 2μm in D,G, H. Panels D-H image DOPE-pLys-647. (I) Simulation snapshots of a simplified lattice model replicating essential elements of the experimental system. Movie is shown in Supplementary Movie 4. (J) The average density of polyelectrolytes as a function of distance from the membrane for the two prewet conditions shown in I. pLys traces show a superposition of pLys tethers and soluble polymers. Beyond the reach of the tethers, the density of both soluble polymers decay exponentially with characteristic length-scales of 0.7 and 1.8 lattice spacings respectively.

We access three distinct regimes in the phase behavior of this coupled system by varying the concentration of soluble pLys and pGlu polymers (Figure 1B). At low concentration, pLys and pGlu polymers remain dispersed in solution and fluorescently labeled pLys-tethers are uniformly distributed on the (dry) membrane. At polyelectrolyte concentrations above saturation (c_sat_), liquid condensates rich in soluble pLys and pGlu polymers are present in solution and can be visualized using a fluorescent pLys conjugate (pLys-532) and by transmitted white light. When condensates contact the bilayer, they wet the membrane, incorporating fluorescently tagged DOPE-pLys tethers (DOPE-pLys-647) at the droplet membrane interface.

At intermediate pLys/pGlu concentrations, below c_sat_ but above another threshold concentration c_prewet_, domains rich in fluorescent pLys-532 and DOPE-pLys-647 tethers assemble at the membrane, but contain fewer pLys-532 probes than wet condensates and are not visible in transmitted light images, indicating they are much thinner than the wavelength of visible light (400 nm). A phase diagram characterizing how c_prewet_ varies as a function of soluble polyelectrolyte and NaCl concentration is shown in Figure 1C. Also shown is the analogous diagram for coexisting condensed and dilute phases in the bulk. Of note, polyelectrolyte rich domains assemble at membranes for pLys/pGlu concentrations close to 20 fold lower than needed for polyelectrolyte condensation in the bulk. The observation that domains are both thin and are stable under conditions where polyelectrolytes remain dispersed in the bulk provide strong evidence that these domains are prewet. Prewet domains of pLys/pGlu polyelectrolytes closely resemble the phase separated domains observed in numerous past studies that coupled purified proteins or nucleic acids to membranes (5–12, 18–21), which are also likely prewet phases.

Circular prewet domains diffuse in two dimensions (2D) within membranes (Supplementary Movie 1) and exhibit behaviors characteristic of 2D liquid phases. Circular domains can coarsen through contact and coalescence (Figure 1D) and can themselves wet one dimensional defects within membrane preparations. Individual DOPE-pLys tethers are mobile but confined within prewet drops, as observed by tracking single molecules over time (Figure 1E) and tabulating the mean squared displacement (Figure 1F). As expected, c_prewet_ varies with tether density (Supplementary Figure 1) and droplet assembly is reversible, both by acutely increasing NaCl and by removing soluble polymers (Figure 1G,H and Supplementary Movies 2,3), although the time-scales vary. The slow melting of drops upon the removal of soluble polymers suggests that the exchange of pLys and pGlu between prewet domains and the 3D bulk is slow, consistent with expectations in this polyelectrolyte system (32). Together, these observations are consistent with prewet domains being fluid surface phases at thermodynamic equilibrium.

Prewetting and wetting of polyelectrolytes can also be demonstrated through simulations of membranes with mobile tethers and two types of soluble polymers (Figure 1I), modifying a simulation scheme reported previously (22) and as described in Methods. Briefly, simulations include mobile, rigid pLys tethers that are confined to the membrane plane and flexible pLys and pGlu polymers free to move in the bulk. Tethers represent a fixed fraction of membrane components (4%) and soluble polymers are held at fixed chemical potential when only surface phases are detected, or at fixed concentration when wetting is observed to retain a finite sized drop. Tethers remain dispersed at a low soluble polymer chemical potential, but are clustered through interactions with soluble polymers at intermediate chemical potentials. Wetting occurs only at high soluble polymer activity when condensed polymer phases are also stable in the bulk. In the intermediate regime, the thickness of the prewet layer is quantified by averaging over domains as described in methods (Figure 1J). Beyond the tethers, which extend a fixed distance from the membrane plane, the density of soluble polymers decays exponentially into the bulk. The decay remains on molecular length-scales in the prewetting regime and becomes macroscopic at the wetting transition (Supplementary Figure S2).

### Proximity to the membrane phase transition potentiates prewetting

We next interrogated prewetting on more compositionally complex membranes of DOPC, DPPC, and cholesterol (Chol) following the experimental scheme represented in Figure 2A. Planar supported membranes constructed from these lipids can demix into two coexisting phases, called liquid-ordered (Lo) and liquid-disordered (Ld) (33). In the absence of soluble polymers, membranes of 36% DOPC, 40% DPPC, 20% Chol, 4% DOPE-pLys-647, and 0.5% DiO are in a single liquid phase at 40°C but contain coexisting Lo and Ld phases at 35°C (Figure 2B). Lipid demixing is detected by imaging the localization of the Ld favoring probe DiO and is fully reversible upon changes in temperature (Supplementary Movie 4). Fluorescent DOPE-pLys-647 tethers partition into the Ld phase.

**Figure 2:**
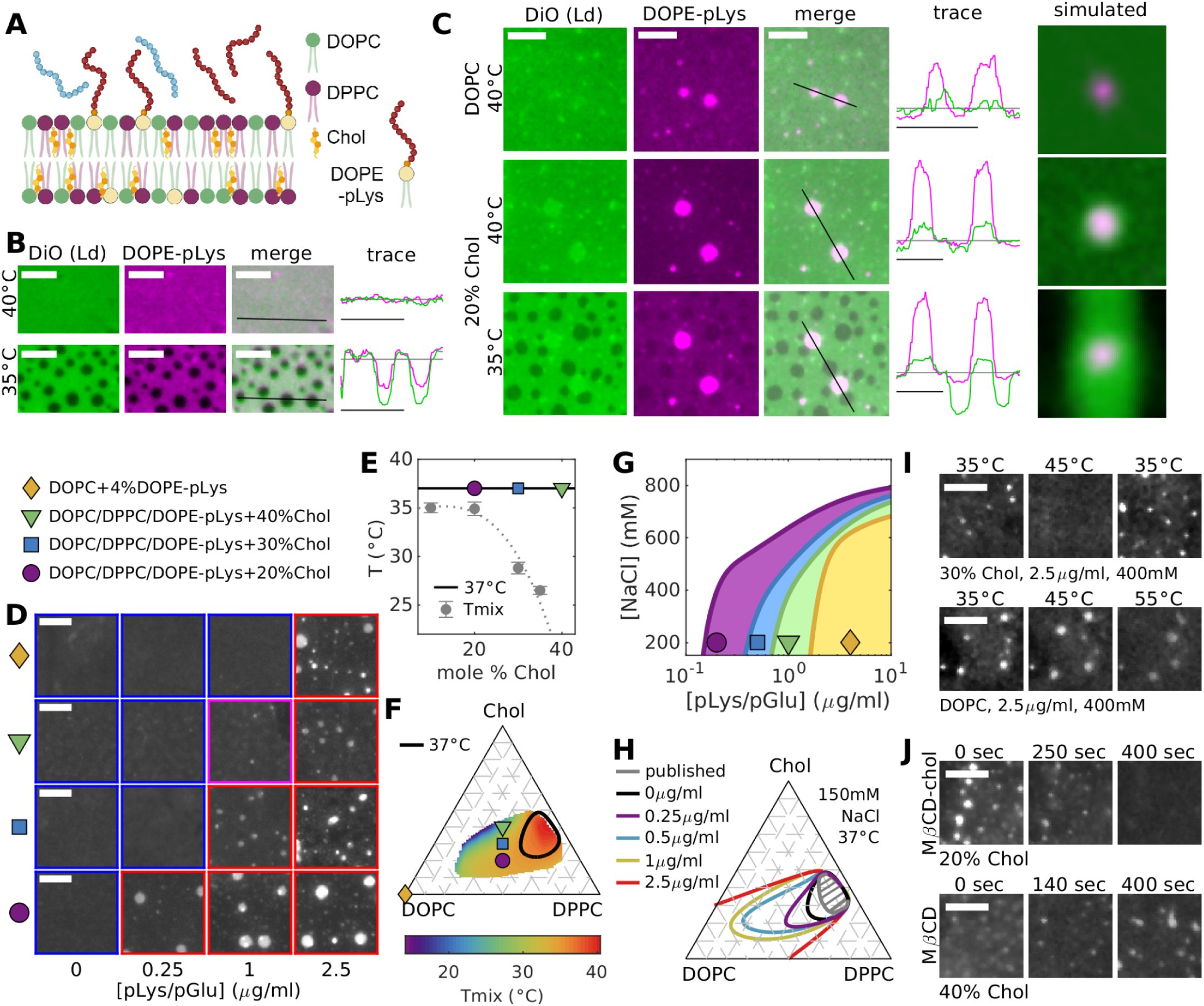
Proximity to the membrane phase transition potentiates prewetting. (A) Schematic representation of the experimental system containing bulk pLys and pGlu in contact with membranes of DOPC, DPPC, Chol, and DOPE-pLys tethers. (B) Membranes of 36%DOPC/40%DPPC/4% DOPE-pLys/20%Chol are in a single phase at 40°C and separate into two coexisting surface phases (Lo and Ld) at 35°C. Both the lipid probe (DiO) and the tether (DOPE-pLys-647) label the Ld phase. (C) Membranes with 2 (top and middle) or 3 (bottom) coexisting phases in experiment (left 4 panels) and simulation (right panels). Experimental systems consist of 96%DOPC/4%DOPE-pLys (DOPC) or 36%DOPC/40%DPPC/4% DOPE-pLys/20%Chol (20% Chol) membranes with 5μg/ml 1:1 w/w pLys/pGlu and 400mM NaCl in solution. Coexisting tether-rich prewet and tether-poor dry phases coexist for both membrane compositions at 40°C with prewet domains also enriching the DiO lipid probe for the 20% Chol membrane. Prewet domains coexist with 2 distinct dry phases at 35°C each with different compositions of labeled tethers and DiO. Traces shown in B,C indicate intensities along the black line normalized by the median value over the entire image for each color channel. Time-averaged densities of tethers and Ld lipids from simulations mimicking experimental conditions. Additional simulation results are shown in Supplementary Figure S2. (D) Representative micrographs of DOPE-pLys-647 in the presence of 1:1 (w/w) mixtures of soluble pLys and pGlu on membranes imaged at 37°C with 400mM NaCl. Distinct membrane compositions are labeled by filled symbols and indicated in the legend which applies to parts D-G. (E) Transition temperatures (Tmix) measured for bare planar membranes without polyelectrolytes. Gray points show the average and standard error over at least 3 measurements. The dotted line is a polynomial fit to guide the eye. (F) Surface depicting the onset of coexisting Lo and Ld phases in bare membranes of DOPC, DPPC, and Chol reproduced from past work (31) along with compositions probed in this study. The black line represents an estimate of the phase boundary for membranes containing 4 mole% DOPE-pLys at 37°C. (G) Phase diagram for membrane compositions depicted in D-F as a function of [NaCl] and soluble [pLys/pGlu]. Individual points are shown in Supplementary Figure 1. (H) Estimated ternary phase boundary for varying soluble pLys/pGlu at T=37°C and [NaCl]=150mM. Phase diagram showing points and images are in Supplementary Figure S5. (I) Prewet domains reversibly dissipate when temperature is raised and lowered in 30% Chol membranes but not in majority DOPC membranes. (J) Prewet domains dissipate or form when Chol is added or removed with MβCD-Chol or MβCD respectively. Images I,J represent frames from movies supplied as Supplementary Movies 5,6. All scale bars are 5μm.

Similar to the majority DOPC membrane of Figure 1, prewet domains form with the addition of soluble pLys and pGlu to this more compositionally complex membrane (Figure 2C). At 40°C, two surface phases are detected. One is a prewet phase rich in DOPE-pLys-647 tethers and soluble polyelectrolytes and the second is a dry phase where tethers are depleted. Prewet domains in this more complex lipid mixture also enrich the fluorescent lipid analog DiO indicating that the two surface phases have distinct compositions of lipid components, distinguishing them from prewet domains in majority DOPC membranes. At 35°C, three surface phases are detected. Again, the prewet phase is enriched in tethers, soluble polyelectrolytes and the Ld marker DiO. Additionally there are two dry phases assigned as Ld-dry and Lo-dry based on the partitioning of the Ld probe DiO. Simulations of complex membranes with mobile tethers qualitatively replicates the phase separation behaviors observed experimentally: coexisting prewet and dry phases can have distinct lipid compositions, prewet domains remain molecularly thin over a wide range of conditions, and three distinct surface phases can coexist at equilibrium (Figure 2C and Supplementary Figure 2). This simulation represents a minor modification of a model we probed extensively in past work, where three coexisting phases were detected over a range of conditions (22). These observations are also in good agreement with past experimental studies that report two (11, 21) or three (11) coexisting surface phases in select compositions of membranes and reconstituted proteins.

A major prediction of our past theoretical work was that proximity to the membrane phase transition potentiates prewetting (22). This effect is experimentally validated by titrating soluble pLys/pGlu on membranes with distinct lipid compositions at fixed temperature (37°C; Figure 2D). We examined 3 Chol containing membranes with miscibility transition temperatures (Tmix) below 37°C. Each membrane contains a varying Chol mole% but retains a fixed ratio of unsaturated (DOPE-pLys and DOPC) to DPPC lipids where DOPE-pLys is held constant at 4 mole% of membrane lipids. In the absence of soluble polyelectrolytes, all membranes are in a single fluid phase at 37°C but are different distances from Ld-Lo coexistence, both in temperature (Figure 2E) and in composition (Figure 2F). Bare membranes with 20% Chol are positioned very close to Tmix (ΔT = 1.1±0.7°C), while bare membranes with 30% or 40% Chol are positioned further from Tmix (ΔT = 8.2±0.6 and >15°C respectively). Prewetting is potentiated in all three membranes as compared to the majority DOPC membrane of Figure 1, with the largest effect in the membrane poised closest to its membrane transition (20% Chol). Chol levels within mammalian plasma membranes also vary between 20 and 40%, depending on cell type and growth conditions, but otherwise have very different lipid compositions than the model membranes employed here (34). Here we vary Chol levels in this range to explore how ΔT impacts the prewetting transition.

Phase diagrams characterizing regions of prewet/dry coexistence as a function of soluble polyelectrolyte and NaCl concentration are shown in Figure 2G and Supplementary Figure S3. Notably, c_prewet_ is 10-fold lower on the membrane containing 20% Chol than on the majority DOPC membrane and prewet domains are present to higher NaCl concentrations. In addition to potentiating c_prewet_, membrane composition influences the adsorption of soluble pGlu to the membrane in both experiment and simulation (Supplementary Figure S4). A more exhaustive exploration of lipid ratios at fixed NaCl concentration (Figure 2H and Supplementary Figure S5) reveals that all compositions with a reported Lo-Ld phase transition potentiate prewetting (compare colored surface in Figure 2F to 1μg/ml contour in Figure 2H). This dramatic and robust potentiation of the prewetting transition provides experimental support for theoretical predictions (22), and extends past experimental work in analogous systems (11, 21). Specifically, this work identifies an extended regime where prewetting depends strongly on membrane-mediated interactions in addition to electrostatic interactions between polyelectrolytes.

For the measurements of Figure 2F-H, prewetting is initiated by titrating soluble pLys/pGlu within the bulk above membranes of fixed composition and temperature. Prewetting can also be reversibly initiated by varying temperature in DOPC/DOPE-pLys/DPPC/Chol membranes, which impacts proximity to the membrane phase transition (ΔT = T-Tmix) and also the magnitude of polymer interactions relative to the thermal energy k_B_T (Figure 2I). Similarly, prewetting can be reversibly initiated at fixed temperature and pLys/pGlu concentration by changing membrane Chol content using methyl-β-cyclodextrin (MβCD; Figure 2J, Supplementary Movies 5,6). Extracting Chol from a 40% Chol membrane decreases ΔT and prewet domains assemble over time, while loading Chol into a 20% Chol membrane increases ΔT and prewet domains melt over time. In these examples, tether surface density (in the case of changing temperature) or tether mole % (in the case of adding and removing Chol) are not held constant. We anticipate that these changes impact but do not dominate the observed effects because c_prewet_ is weakly dependent on tether density in DOPC membranes in the vicinity of 4 mole% DOPE-pLys (Supplementary Figure 1).

### Protein prewetting at the plasma membrane is tuned by membrane properties

The same prewetting behaviors detected in the model system can be reconstituted within cells using existing optogenetic tools (35). We use a 3 component modular system consisting of a Cry2-motif conjugated to an anti-GFP nanobody (Cry2-ɑGFP), a soluble CIB1 protein conjugated to a multimeric protein (CIB1-MP), and a GFP anchored to the plasma membrane through the palmitoylated and myristoylated membrane binding domain of Lyn kinase (PM-GFP). Each protein component is conjugated to a distinguishable fluorescent tag. Upon exposure to blue light, Cry2-ɑGFP undergoes a conformational change that enables binding to CIB1-MP and other activated Cry2 molecules, as represented schematically in Figure 3A.

**Figure 3:**
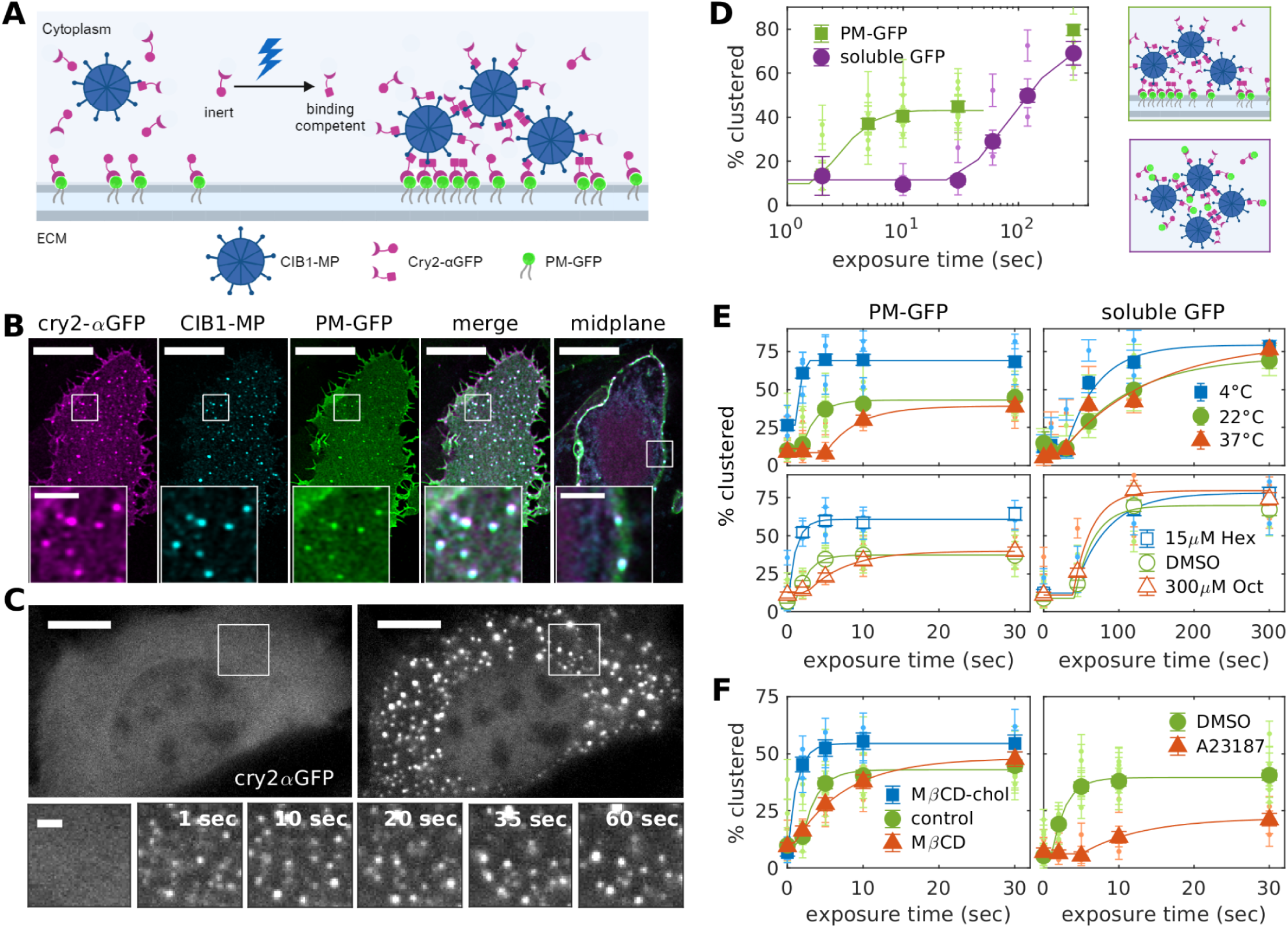
Prewetting at the plasma membrane is sensitive to perturbations of membrane structure and composition. (A) Schematic representation of the optogenetic system containing multivalent C1B1-MP, Cry2-αGFP, and a membrane anchored PM-GFP. Upon exposure to blue light, Cry2-αGFP can interact with itself and/or CIB1-MP. (B) Images of the 3 components co-expressed in U2OS cells chemically fixed after 5 sec of light exposure. Puncta enriched in all 3 components are present on the dorsal surface and localize to the plasma membrane in a midplane slice. (C) (Top) Live U2OS cell imaged at 37°C before (left) and after a brief exposure to blue light. (Bottom) Insets from above showing domain coarsening over time. Images are frames from Supplementary Movie 7. (D) Curves representing the percentages of cells exhibiting Cry2-αGFP/CIB1-MP puncta in fields of cells chemically fixed after the varying exposure times to blue light. Cells either co-express PM-GFP or a soluble form of GFP. Representative fields for several light exposures are shown in Supplementary Figure S8. (E) Curves representing the percentage of cells with Cry2-αGFP/CIB1-MP puncta in cells treated and chemically fixed at different ambient temperatures (top) or in the presence of n-alcohols (bottom). (F) Curves representing the percentage of cells with Cry2-αGFP/CIB1-MP puncta when treated and chemically fixed after Chol modulation with MβCD or MβCD-Chol (left) or in the presence of the calcium ionophore A23187 (right). Analogous curves for cells expressing soluble GFP are in Supplementary Figure S12. In B and C, scale-bars are 10 μm in main images and 2 μm in insets. Large points and error bars in D-F represent a variance weighted mean and error of the weighted mean over biological replicates shown as small symbols with errors derived from counting statistics as described in Methods. Numbers of replicates for each dataset are listed in Supplemental Table 2. Lines in D-F are fits to an exponential function with a lag, as described in Methods. Movies showing perturbations of E,F applied in live cells are shown in Supplementary Movies 9-12.

When transiently co-expressed in U2OS cells, PM-GFP uniformly labels the plasma membrane, Cry2-ɑGFP labels the membrane and is also abundant in the cytoplasm and nucleus, and CIB1-MP is cytoplasmic and sometimes labels internal structures (Supplementary Figure S6). Upon exposure to blue light, Cry2-ɑGFP, CIB1-MP, and PM-GFP co-assemble into puncta exclusively observed at the plasma membrane when imaged by Ariryscan confocal microscopy (Figure 3B and Supplementary Figure S7). Imaging only Cry2-ɑGFP in live cells reveals that circular domains are mobile and coarsen through coalescence (Figure 3C and Supplementary Movie 7), resembling the phase separated prewet domains observed in the fully reconstituted system of Figures 1-2. In the absence of blue illumination, domains melt over a time-scale of roughly 10 minutes at room temperature as Cry2 relaxes into its low-affinity state (Supplementary Movie 8).

We next quantified the stability of membrane bound Cry2/CIB1-MP puncta by systematically varying exposure to blue light at room temperature (22°C), effectively titrating the concentration of binding competent Cry2-ɑGFP. Briefly, we quantify the fraction of cells containing Cry2ɑ-GFP/CIB1-MP puncta within a population of cells transiently expressing Cry2-ɑGFP, CIB1-MP, and plasma membrane targeted PM-GFP. Cells from the same transfection were plated onto separate dishes overnight, then exposed to different durations of blue light prior to chemical fixation and imaging, as described in Methods and Supplementary Figure S8. We note that the contrast of PM-GFP within Cry2-ɑGFP/CIB1-MP puncta is often subtle, which we attribute to their being only a small fraction of activated Cry2-ɑGFP bound to GFP when cells are exposed to a short duration of blue light. Results are summarized in the phase diagram of Figure 3D, which show that much shorter light exposures are required to form Cry2-ɑGFP/CIB1-MP puncta in cells expressing PM-GFP as compared to cells expressing a soluble monomeric GFP that also binds to Cry2-ɑGFP but does not contribute to clustering. Our observation is in good agreement with past findings with these constructs (36) and applies even though expression of Cry2-ɑGFP and CIB1-MP are comparable in the PM-GFP and GFP containing samples (Supplementary Figure S9). Plasma membrane associated Cry2-ɑGFP/CIB1-MP puncta extend into cell interior when cells cells express PM-GFP tethers are chemically fixed after after a long exposure to blue light (300s), closely resembling Cry2-ɑGFP/CIB1-MP puncta observed in cells expressing soluble GFP (Supplementary Figure S7). The titration of Figure 3D indicates that the fraction of cells expressing PM-GFP saturates for exposure times less than 30s, then increases with a long exposure (5 min), matching the value observed for cells expressing soluble GFP. We attribute the smaller initial saturation value to there being a requirement for individual cells to express sufficient levels of PM-GFP, Cry2-ɑGFP, and CIB1-MP to assemble puncta at short light exposures, while individual cells need only express Cry2-ɑGFP at sufficiently high levels to assemble puncta at long light exposures (Supplementary Figure S9).

We have previously argued that mammalian cell plasma membranes are in a single phase but near a membrane miscibility transition (27, 37), and we next sought to investigate if perturbations of plasma membrane phase behavior alter the stability of plasma membrane localized Cry2-ɑGFP/CIB1-MP puncta in U2OS cells (Figure 3E). We first varied the ambient temperature of cells during light exposure and chemical fixation. Puncta assembly was potentiated when temperature was lowered to 4°C, meaning that more cells contained puncta when exposed to shorter light exposures than in cells held at 22°C, and more cells contained puncta at saturating light doses at 4° vs. 22°C. In contrast, puncta assembly was inhibited by elevating temperature to 37°C, meaning that fewer cells contained puncta at short light exposures than cells held at 22°C. This temperature effect was also evident in a representative live cell (Supplemental Movie 9). In this cell, Cry2-ɑGFP remains uniformly distributed after a short exposure to blue light at 37°C, but condenses into puncta following a temperature quench to 15°C. Changes in ambient temperature alter proximity to Tmix, but also impact Cry2 deactivation kinetics and the relative magnitude of Cry2/Cry2 and Cry2/CIB1-MP interactions compared to the thermal energy k_B_T. These non-membrane effects are apparent when interrogating the light exposure time dependence of cytoplasmic Cry2/CIB1-MP puncta in cells expressing Cry2-ɑGFP, CIB1-MP, and soluble GFP (Figure 3E). In this case, lowering temperature to 4°C modestly potentiates cytoplasmic Cry2/CIB1-MP puncta assembly when compared to cells investigated at 22°C.

The specific impact of the membrane is better isolated using chemical perturbations of the membrane transition at fixed ambient temperature. To accomplish this, we used two n-alcohols shown previously to modulate transition temperatures in isolated giant plasma membrane vesicles (GPMVs) and to alter contrast of domains in live B cell membranes (37, 38). GPMVs retain many properties of the cell membranes from which they are derived, but also differ in important ways, including exhibiting macroscopic phase separation under conditions where intact cell membranes remain in a single phase (39). GPMVs isolated from U2OS cells separate into coexisting liquid phases below room temperature, and Tmix increases in the presence of 15 μM hexadecanol and decreases in the presence of 300 μM octanol (Supplementary Figure S10). Pretreating intact U2OS cells expressing Cry2-ɑGFP, CIB1-MP, and PM-GFP with 15 μM hexadecanol potentiated puncta assembly, producing a curve analogous to the one observed when ambient temperature was lowered to reduce ΔT. In contrast, pretreating intact U2OS cells with 300 μM octanol inhibited puncta assembly, producing a curve analogous to the one observed when ambient temperature was raised to increase ΔT. The melting of domains and desorption of Cry2-ɑGFP from the plasma membrane can also be detected in live cells after acute treatment with 300 μM octanol at fixed ambient temperature (Supplemental Movie 10 and Supplementary Figure S11). Unlike changing ambient temperature, octanol and hexadecanol treatments do not significantly impact the assembly of cytoplasmic puncta over a wide range of light exposure durations. This supports the conclusion that n-alcohol treatment alters the stability of Cry2/CIB1-MP puncta via its impact on membrane properties, likely through their effect on Tmix.

We next explored two additional perturbations that alter membrane properties without directly altering the magnitude of protein interactions (Figure 3F). The first involved perturbing plasma membrane Chol, where we found that supplementing plasma membrane Chol with MβCD-Chol stabilized Cry2/CIB1-MP puncta and depleting Chol with MβCD destabilized Cry2/CIB1-MP puncta in both chemically fixed and live cells (Supplementary movies 11,12). These MβCD treatments represent large perturbations of cellular Chol content (as much as ±50% (40, 41)) and can alter the concentration or localization of membrane components besides plasma membrane Chol (42, 43). We note that the impact of Chol changes in cells is inverse to that observed in the model membranes of Figure 2, and attribute this to the significant difference in lipid composition between models and cells. As expected, MβCD treatment does not alter the light sensitivity of Cry2-ɑGFP/CIB1-MP puncta assembled in the presence of soluble GFP (Supplementary Figure S12) indicating that these treatments do not directly alter Cry2/Cry2 or Cry2/CIB1-MP interactions. As a final perturbation, we treated cells with the calcium ionophore A23187. Elevated cytoplasmic Ca^2+^ activates the plasma membrane localized scramblase TMEM16, relaxing some plasma membrane leaflet asymmetry (43, 44) as detected through the binding of Annexin V on the extracellular leaf (Supplementary Figure S13). Cells pretreated with A23187 exhibit greatly diminished Cry2/CIB1-MP puncta even after long light exposure, and acute A23187 treatment leads to the abrupt dissociation of domains in live cells (Supplementary Movie 13). Many processes at the plasma membrane and within the cytoplasm are activated by high cytoplasmic Ca^2+^ (45, 46), leading to many possible off-target effects. Even so, A23187 pre-treatment does not greatly impact PM-GFP levels at the plasma membrane (Supplementary Figure 11), or Cry2-ɑGFP/CIB1-MP puncta assembled in the presence of soluble GFP (Supplementary Figure S12), indicating that relevant effects are limited to those that impact the plasma membrane.

The measurements described above provide strong evidence for protein prewetting in a cellular context. First, expression of the PM-GFP tether is required to detect puncta at short light exposures in cells coexpressing Cry2-ɑGFP and CIB1-MP (Figure 3D). Second, the light exposure duration required to assemble Cry2-ɑGFP/CIB1-MP puncta changes in the presence of perturbations which act exclusively at the membrane surface (Figures 3E,F). These two observations are consistent with a prewetting transition and inconsistent with wetting, where the stability of domains in the thermodynamic limit is determined only by interactions in the bulk (23). Furthermore, puncta are only observed at the plasma membrane by Airyscan confocal microscopy (Figure 3B and Supplementary Figure S7) indicating that they are thinner than the resolution limit of the microscope (<350nm). Overall, puncta observed in cells are highly analogous to the prewet domains detected in model membranes with a nearby miscibility transition as described in Figure 2.

### Protein prewetting at the ER membrane is potentiated by perturbations of ER lipids

Protein prewetting at the cytoplasmic leaflet of the ER can be interrogated using the same optogenetic tools, by replacing PM-GFP with Sec61-GFP (Figure 4A), a transmembrane member of the translocon machinery often used as a general ER marker. Sec61-GFP that is initially uniformly distributed across the ER (Supplementary Figure 14) co-clusters with Cry2-ɑGFP and CIB1-MP after exposure to blue light (Figure 4B). Live cell imaging of only Cry2-ɑGFP reveal dynamic puncta that also coarsen over time (Figure 4C and Supplementary Movie 14). Profiling expression levels by flow cytometry reveals that while Sec61-GFP is expressed at a much lower level than PM-GFP, while Cry2-ɑGFP and CIB1-MP expression is comparable to those observed in measurements described in Figure 3 (Supplemental Figure S15). Even so, puncta containing these proteins assemble in the presence of the Sec61-GFP tether under conditions where they remain dispersed without membrane tethering (Figure 4D), consistent with domains induced by short light exposures being prewet.

**Figure 4:**
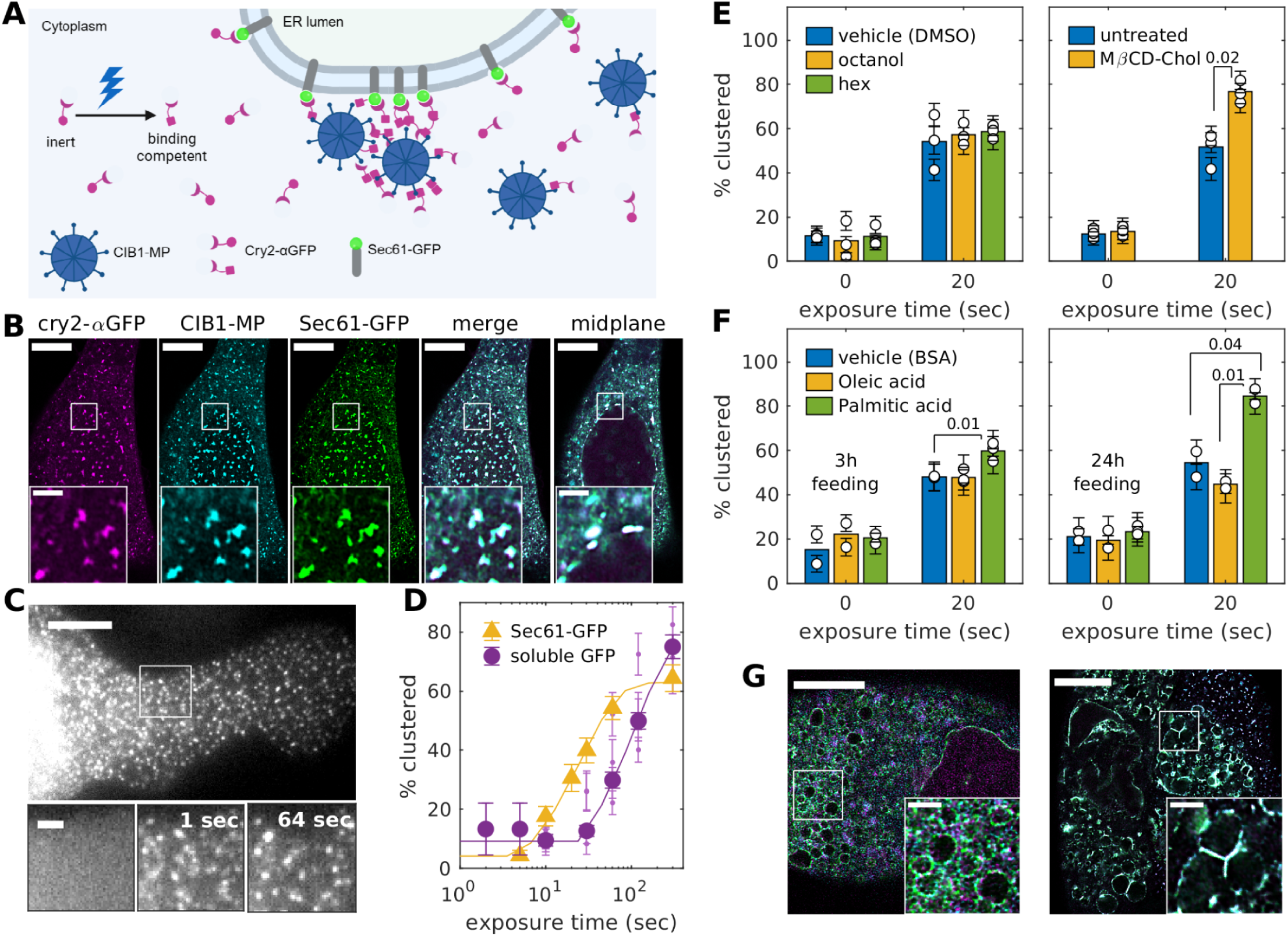
Prewetting at the ER cytoplasmic leaflet is potentiated by perturbations of membrane composition. (A) Schematic representation of the optogenetic system containing multivalent C1B1-MP, Cry2-αGFP, and Sec61-GFP localized to the ER membrane. (B) Airyscan confocal images of the 3 components co-expressed in U2OS cells chemically fixed after 10 sec of light exposure at 22°C. Puncta enriched in all 3 components are present throughout the cell and larger domains are seen proximal to the nuclear envelope. (C) (Top) Cry2-αGFP-SNAP-SiR imaged in a live U2OS cell at 37°C after a brief exposure to blue light. (Bottom) Insets showing domain coarsening over time. Images are frames from Supplementary Movie 14. (D) Curves representing the percentages of cells exhibiting Sec61-GFP/Cry2-αGFP/CIB1-MP puncta in fields of cells chemically fixed after the varying exposure times to blue light at 22°C. Points representing soluble GFP are replotted from Figure 3D. (E) Bars representing the percentage of cells with Cry2-αGFP/CIB1-MP puncta in cells treated and chemically fixed in the presence of n-alcohols (left) or after Chol loading with MβCD-Chol (right). (F) Bars representing the percentage of cells with Cry2-αGFP/CIB1-MP puncta in cells chemically fixed after 3 h (left) or 24 h (right) of fatty acid feeding in culture media at 37°C. (G) Airyscan confocal images of spherical Sec61-GFP labeled ER structures observed in a subset of PA treated cells chemically fixed without (left) or with (right) 20 s of blue light exposure. Scale-bars are 10 μm in main images and 2 μm in insets. Large points and error bars in D-F represent a variance weighted mean and error of the weighted mean over independent replicates, shown as small symbols with errors derived from counting statistics as described in Methods. Numbers of replicates for each dataset are listed in Supplemental Table 2. p-values shown in E,F are tabulated using unpaired T-tests from two or three biological replicates. Lines in D are fits to an exponential function with a lag, as described in Methods. Representative fields for several conditions are shown in Supplementary Figure S15.

We next explored if perturbations of Cry2-ɑGFP/CIB1-MP prewetting at the plasma membrane also impacted the stability of Cry2-ɑGFP/CIB1-MP domains at the ER, focusing on an illumination time near the midpoint of the titration (20 sec). Unlike at the plasma membrane, octanol and hexadecanol had no impact on prewetting at the ER, although Chol loading through MDCD-Chol potentiated ER prewetting (Figure 4E). In live cells, prewet domains assembled within seconds after the addition of MDCD-Chol (Supplementary Movie 15). Chol manipulation induces endogenous contact sites between the ER and plasma membrane (47), and we speculate that this could be due, at least in part, to membrane mediated effects. Past work also indicates that ER membrane composition can be altered by supplementing growth media with fatty acids, with palmitic acid supplementation acting to rigidify ER membranes while oleic acid supplementation does not (48). More cells contain prewet domains after 20s of light exposure when U2OS cells incubated with palmitic acid for 3 or 24 h than in cells treated for the same times with either oleic acid or a BSA carrier control (Figure 4F).

Past work indicates that prolonged incubation with palmitic acid induces ER stress. Consistent with this we find that some cells exhibit the phenotype shown in Figure 4G in which Sec61-GFP labeled membranes swell and take on spherical shapes, allowing for the direct visualization of isolated ER membranes by confocal imaging. Similar to prewet domains at the plasma membrane, Cry2-ɑGFP and CIB1-MP proteins colocalize at the surface of ER spheres in cells chemically fixed after exposure to blue light, consistent with domains being prewet. Images also suggest that prewetting can facilitate adhesion between adjacent membranes. In cells chemically fixed without blue light exposure, spherical Sec61-GFP labeled structures appear distinct, even when closely packed within the cell interior. In contrast, cells exposed to blue light adhere, sharing boundaries rich in Cry2-ɑGFP and CIB1-MP.

### Contact sites driven by prewetting are sensitive to membrane composition

We next sought to directly explore if prewetting could drive membrane adhesion in our reconstituted system, using large unilamellar vesicles (LUVs) containing DOPC and DOPE-pLys tethers and soluble pLys and pGlu polyelectrolytes by dynamic light scattering (DLS), a method that reports on the size distribution of diffusing objects in suspension (Figure 5A). Here we found that the size distribution of vesicles both broadened and shifted to larger average values near c_prewet_ as determined in the planar bilayer measurements of Figures 1 and 2, consistent with vesicles adhering into small aggregates. To support this conclusion, we conducted fluorescence cross-correlation spectroscopy (FCCS) measurements to measure the co-diffusion of differently colored LUVs made of DOPC and DOPE-pLys tethers (Figure 5B). Here we found a cross-correlation between DiI and DiO labeled LUVs only when pLys and pGlu polyelectrolytes were included at concentrations above c_prewet_. In addition, both DLS and FCCS measurements indicate that vesicle aggregation occurs at lower soluble polyelectrolyte concentrations when LUVs are composed of a fixed ratio of unsaturated (DOPE-pLys and DOPC) and DPPC lipids and 20% Chol, indicating that the avidity of this adhesive reaction is sensitive to membrane lipid content, consistent with c_prewet_ being lower for this membrane composition (Figure 5C).

**Figure 5:**
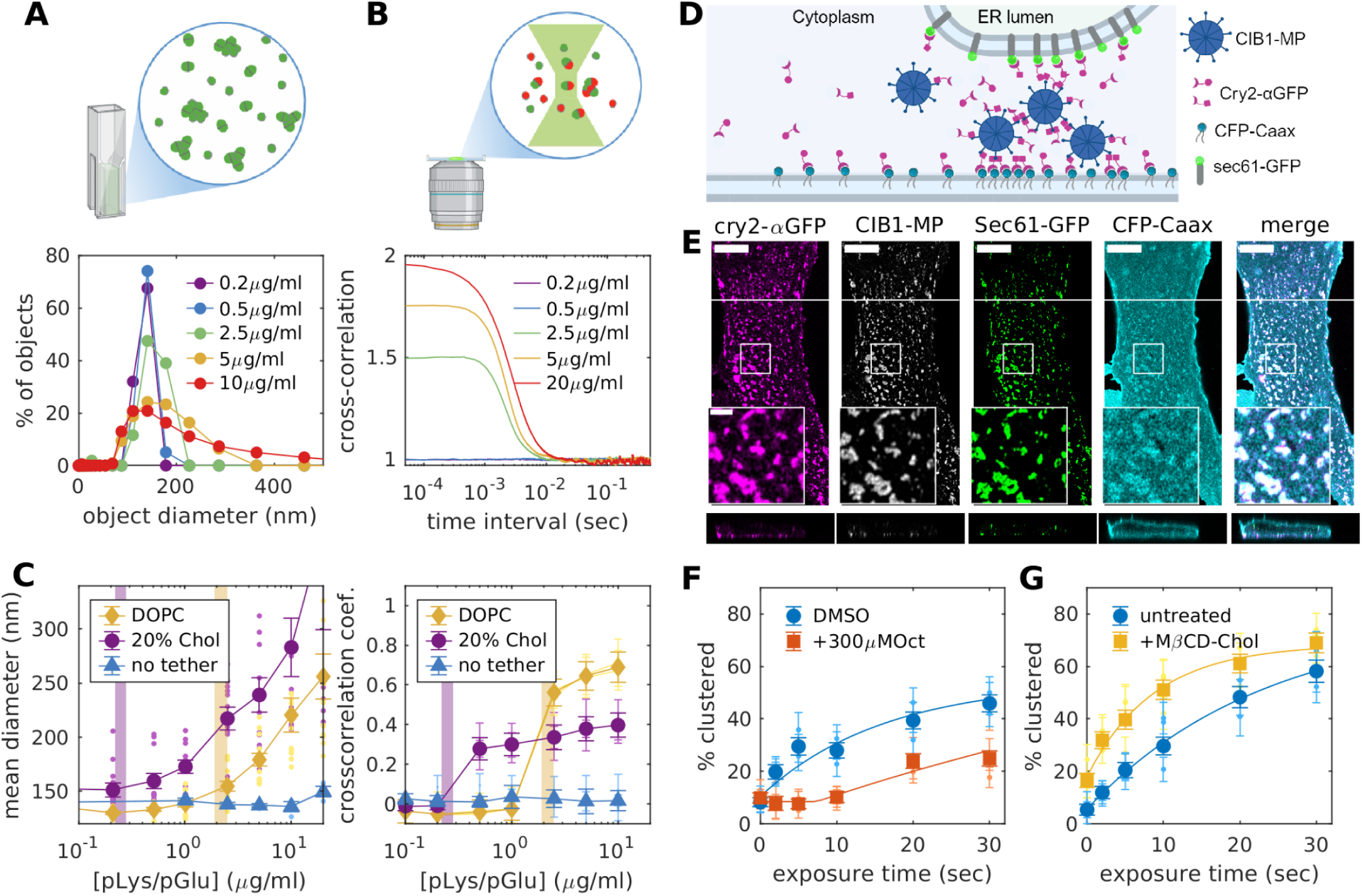
prewetting facilitates membrane adhesion in models and cells. (A) LUV aggregation is observed through DLS measurements with the addition of 1:1 w/w mixtures of soluble pLys and pGlu as a broadening and an upwards shift in histograms representing the distribution of sizes of diffusing objects. (B) LUV aggregation is detected by FCCS as an increase in the cross-correlation amplitude between differently colored vesicles. In A,B vesicles contain 96% DOPC and 4% DOPE-pLys and aggregation is detected after the addition 2.5μg/ml of a 1:1 w/w mixture of soluble pLys and pGlu. (C) Summary of DLS (left) and FCCS (right) results over a range of soluble polymer concentrations for LUVs of 96% DOPC + 4% DOPE-pLys and 36% DOPC + 4% DOPE-pLys + 40% DPPC + 20% Chol tabulated as described in Methods. Vertical bars correspond to c_prewet_ for each membrane mixture. The no tether control contains DOPE-DBCO lipids not conjugated to N_3_-pLys. The mean diameter represents the centroid of the diameter distribution in the DLS measurement and the cross-correlation coefficient is 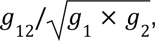 where *g*_12_ is the amplitude of the cross-correlation curve and *g*_1_and *g*_2_are amplitudes of each autocorrelation curve in the FCCS measurement. (D) Schematic of the optogenetic system containing CIB1-MP, Cry2-αGFP coexpressed with CFP-Caax that labels the inner leaflet of the plasma membrane and Sec61-GFP that labels the ER. Cry2-αGFP binds to both GFP and CFP. (E) Images of the 4 components at the ventral surface (top) of a U2OS cell chemically fixed after 5 sec of blue light exposure. Puncta are enriched in all 4 components and are most prominent at the cell periphery (bottom, x-z slice at white line). CIB1-MP is not included in the merge and scale-bars are 10μM in the main panel and 2μm in the inset. (F,G) Curves representing the percentages of cells exhibiting Cry2-αGFP/CIB1-MP puncta as a function of light exposure in fields of cells expressing Cry2-αGFP,CIB1-MP, Sec61-GFP and CFP-Caax in the presence of the indicated pre-treatments. Large points and error bars in C,F,G represent means and SEMs over independent measurements, shown as small symbols with errors if available. Numbers of replicates for each dataset are listed in Supplemental Table 2. Lines in F,G are fits to an exponential function with a lag, as described in Methods.

In the cellular system, we further explored prewetting induced membrane adhesion by generating light-induced contact sites between the plasma and ER membranes using U2OS cells expressing Cry2-αGFP and CIB1-MP along with distinguishable tethers at the plasma membrane (CFP-Caax) and endoplasmic reticulum (Sec61-gfp) that both engage with the αGFP nanobody (Figure 5D). In the absence of blue light exposure, Sec61-GFP and CFP-Caax label the ER and plasma membrane respectively, CIB1-MP is cytosolic, and Cry2-αGFP labels membranes but is predominantly in the cytoplasm (Supplementary Figure S16). Cells exposed to short doses of blue light contain puncta labeled by all 4 expressed proteins that localize exclusively to the cell periphery (Figure 5E). Under these conditions, Sec61-GFP can also be visualized under total internal reflection illumination indicating that extended regions of the ER membrane reside within several hundred nm of the plasma membrane (Supplementary Figure S17). Similar to prewet domains at individual membranes, light induced contact sites can be potentiated or inhibited through perturbations that partition exclusively into membranes (Figure 5 F,G).

While all Cry2-αGFP/CIB1-MP puncta appear to enrich both Sec61-GFP and CFP-Caax upon low light exposure, longer exposure times produce cells that also contain puncta with only one membrane tether (Supplementary Figure S16). This suggests that puncta formed upon short exposures to blue light assemble cooperatively. Consistent with this conclusion, Sec61-GFP puncta assemble at lower light exposures in cells expressing Cry2-αGFP, CIB1-MP, and CFP-Caax than in cells expressing Sec61-GFP, Cry2-αGFP, and CIB1-MP, even though the expression levels of these three proteins are reduced in cells also expressing CFP-Caax (Supplemental Figure S18). A similar result is obtained in our reconstituted system, in which LUVs containing DOPC and DOPE-pLys tethers co-diffuse with Chol containing LUVs at soluble polymer concentrations below c_prewet_. (Supplemental Figure S19) These findings suggest that coupling distinct membranes via soluble proteins can potentiate prewetting, although additional work is needed to decipher detailed mechanisms in this more complex system.

## DISCUSSION

Here we have demonstrated that membranes and bulk components prone to liquid-liquid phase separation can form surface phases through prewetting. We found that these prewet domains are exquisitely sensitive to properties of membranes through modeling and experiments in controlled reconstituted systems and cells. In reconstitution, we coupled two systems with well characterized miscibility transitions (29, 33), and found that prewet polyelectrolyte domains formed well outside the coexistence region of either transition. Moreover, we found that tuning membrane lipids could reduce the concentration of soluble polymer required to observe prewet domains, c_prewet_ by more than an order of magnitude. In cells, we exploited available optogenetic tools (35) to generate prewet domains at the plasma membrane inner leaflet and on the cytoplasmic leaflet of the ER. Here too we found that the stability of prewet domains depends strongly on membrane composition, probed through a range of membrane perturbations. Some of these perturbations have well characterized impacts on the membrane phase transition in isolated and intact plasma membranes (37, 38), supporting a role of the membrane phase transition in tuning the stability of prewet protein assemblies even though both membranes in isolation are uniform liquids. Finally, we demonstrated that prewetting can facilitate the assembly of contact sites whose stability depends strongly on membrane composition. There is remarkable consistency across the models and cell measurements, supporting the view that universal aspects of these phase transitions underlie the assembly of prewet domains in cells.

These studies add to a large body of work interrogating the biophysical and biological roles of condensates at membranes (5–22, 49–52) by implicating both protein and lipid phase transitions in the assembly of membrane domains. Long-standing theories of domains at the plasma membrane assumed that lipid-interactions were sufficient to confine membrane proteins to functional signaling platforms (28, 53), while more modern views posit that signaling platforms assemble exclusively from multivalent protein interactions (54). The current work suggests that cells likely exploit emergent thermodynamic behaviors of both membrane and bulk components to generate these functional assemblies in cells. Also highlighted by this work is the importance of the tethers which couple these two systems. In our reconstituted system tethers consist of a small mole fraction of lipids attached to polyLys polymers, and in our in vitro work we employ Cry2 bound to lipid anchored GFP. But tethers can take on many forms, ranging from cytoplasmic domains of receptors or adapters phosphorylated within signaling cascades (5, 11, 51), membrane proteins targeted for degradation through ubiquitination (55, 56), or even specific phosphatidylinositol lipid species whose levels can change abruptly during a range of cellular events (57, 58). Reactions which locally produce binding sites on a small number of tethers can efficiently seed thermodynamic domains that would require orders of magnitude more modifications of bulk molecules. But activating tethers can still lead soluble components to rapidly condense into domains which resemble bulk condensates, but stable at much lower concentrations. Thus tether production can be an efficient and rapid way to build signaling platforms or scaffold cellular processes.

Coupling biopolymer and membrane phase transitions produces a new transition that exhibits qualities characteristic of each individual system. Biopolymers condense into phases highly enriched in specific components and their stability can be regulated via conformational changes or posttranslational modifications, enabling specific biological functions (59). Membrane lipids can only separate into two distinct liquid phases, but are more easily tuned to compositions with emergent properties that enhance sensitivity, through proximity to the membrane phase transition (27). Coupling these transitions retains the regulated specificity of condensates and the sensitivity of the membrane transition, enabling phase separation even when condensing biopolymers are dilute. Moreover, these domains become exquisitely sensitive to properties of the surface, which includes perturbations that impact the membrane phase transition, even when crossing into its decoupled coexistence region would require large changes in temperature and/or composition

Many proteins can phase separate into liquid domains in purified systems or with over-expression even when they remain dispersed when expressed at endogenous levels within cells (60), leading to questions regarding the relevance of this miscibility transition to cell biology. Similar questions have been raised regarding the miscibility transition present in mammalian cell plasma membranes, which is exclusively observed in perturbed systems away from physiological conditions. Our work suggests that the molecules that make up biomolecular condensates can play important roles even well outside of bulk phase coexistence, through interactions with surfaces or other structures capable of enhancing condensation. For example, while purified components of the postsynaptic density separate into a bona fide bulk liquid phase when reconstituted at high concentrations, their role in vivo seems to be in forming prewet surface phases, which requires far lower concentrations (6). Postsynaptic membranes are also enriched in lipids associated with the Lo phase (61), possibly indicating that cells exploit the membrane transition to enhance their stability.

While most existing work has focused on the presence and properties of phase separated domains themselves, our work indicates that coupling condensate prone biopolymers to surfaces can drive new phenomena. The current study demonstrates that condensing proteins or polyelectrolytes can drive the assembly of membrane contact sites, but it is conceivable that prewetting or related phenomena could drive other types of structural transitions. For example, tethering phase separating bulk components to membranes through curvature sensitive motifs could promote the clustering of synaptic vesicles in neurons (14) and small intracellular vesicles in B cells (62), or the assembly of caveolae (63) and viral envelopes (64, 65), although so far none of these phenomena have been discussed in the context of prewetting transitions. Prewetting and related surface phenomena could apply to other surfaces in cells, such as microtubules (66) or DNA (67). More broadly, we propose that prewetting and related surface phenomena represent an unexplored avenue into the functional roles of biomolecular condensates in cell biology.

## MATERIALS AND METHODS

### Materials

All chemicals were purchased from suppliers and directly used without any further purification. All lipids including 1,2-dioleoyl-sn-glycero-3-phosphocholine (DOPC), 1,2-dipalmitoyl-sn-glycero-3-phosphocholine (DPPC), 1,2-dioleoyl-sn-glycero-3-phosphoethanolamine-N-dibenzocyclooctyl (DOPE-DBCO), and cholesterol (Chol) were purchased from Avanti Polar Lipids (Alabaster, AL USA). MβCD (CAS#128,446-36-6) was purchased from TCI Chemicals (Portland, OR). Poly (l-glutamic acid sodium salt) (pGlu), CAS#26247-70-0 (MW=7500Da), Poly(l-lysine hydrobromide) (pLys), CAS#25988-63-0 (MW=6300Da) and Poly(l-lysine hydrobromide) azide (pLys-N_3_), CAS# 25988-63-0 (MW=6300Da) was purchased from Alamanda polymers (Huntsville, AL, USA).

Fluorescent lipid analog 3,3’-Dioctadecyloxacarbocyanine Perchlorate (DiO-C18) (Cat#: D3898), DiI-C₁₂ (Cat#D383), Alexa Fluor 532 NHS Ester (Succinimidyl Ester) (Cat# A20101MP), Palmitic acid (CAS#57-10-3), Oleic acid (CAS#112-80-1) and Ionophore A23187 (Cat# J63020.MA) were purchased from Thermofisher Scientific (Waltham, MA, USA). CF 647 succinimidyl ester (Cat# SCJ4600048) was purchased from millipore sigma. SNAP-Cell 647-SiR (Cat#S9102S) was purchased from New England Biolabs. Dithiothreitol (DTT) (Lot#SLBX8626), Paraformaldehyde (PFA) (CAS#30525-89-4) and other remaining reagents were purchased from MilliporeSigma (Burlington, MA, USA).

### Fluorescent labeling of polymers

pLys-N3, pGlu, and pLys (1eq) were modified with a fluorophore containing an NHS ester (5eq) in PBS buffer (pH= 7.4) at room temperature for 1 hour. Unreacted probe was removed by passing the reaction mixture through Amicron Ultra column with 3k cutoff size. pLys-N3 was coupled to CF 646 and pLys and pGlu were each conjugated to Alexa 532.

### Preparation of supported lipid bilayers

Lipid bilayers were prepared by spin-coating following previous protocols (17). Briefly, the specified lipid solutions plus an additional 0.2 mole% DiO were measured from concentrated stock solutions in chloroform and dried under vacuum. The dried lipid film was resuspended in 96% chloroform and 4% methanol to a final lipid concentration of 1.5mg/ml, then 40ul of this solution was spin coated on a clean glass substrate for 30 seconds at 3000 rpm. Glass substrates were either conventional coverslips (Thermo Fisher scientific Cat# 174969) or smart glass substrates (18x18x0.17 mm³, Interherence GmbH, Erlangen, Germany) for temperature controlled measurements. In both cases, substrates were plasma cleaned (Harrick plasma PDC-32G) prior to spin-coating. The glass supported lipid film was then dried under vacuum for 30 min to remove residual solvent. The dried lipid layer was rehydrated with 200μl phosphate buffered saline (PBS), prior to labeling with 2μg/ml CF647-pLys-N3 using a copper free click reaction with 18:1 DBCO PE at 37°C temperature for 30 minutes. Unreacted polymer was removed by exchanging the buffer above the membrane three times with PBS or PBS supplemented with NaCl. Prior to additional experimentation, DiO (blue excitation) and DOPE-pLys-647 (red excitation) were imaged to verify pLys conjugation and the presence of a contiguous bilayer.

### Preparation of Large Unilamellar Vesicles

1.5mg of the specified lipid ratios were measured from stock solutions in chloroform then dried under the vacuum. The dried lipid film was resuspended in 1 ml of phosphate buffer with 400mM NaCl at 85°C and the solution was extruded through Avanti mini-extruder (610020-1EA) with a polycarbonate membrane (610005-1EA) with 100 nm pore size to form LUVs. If indicated, LUVs were modified with 1µg/ml poly (L-Lysine Hydrobromide) azide at 65°C for 20 minutes.

### Preparation of polyelectrolyte solutions

Unmodified pLys and pGlu were separately diluted from concentrated stocks into solutions containing phosphate buffer (pH 7.4) and the desired NaCl concentration, then mixed in a 1:1 (w/w) ratio immediately prior to observation, typically within the imaging chamber. In some measurements, 10% of the respective polymer was replaced by its fluorescent analog (pLys-532 or pGlu-532) to enable imaging of soluble components.

### Preparation of MβCD-chol

MβCD (3 mg/ml) was dissolved in a PBS buffer and degassed for 30 minutes under vacuum and stored under Argon to prevent oxidation. Saturated MβCD/Chol solutions were prepared by first wetting 10 mg of powdered cholesterol with 200µL of methanol, drying under nitrogen, then resuspending in 3mg/ml MβCD solution (15 ml). This suspension was vortexed then sonicated (twice each for 2min) using a Branson bath Ultrasonifier (MODEL S450A, Process Equipment & Supply Inc, North Olmsted OH). This suspension was stored at room temperature with continuous rotation and was typically used at least 12 h after preparation. Immediately prior to an experiment, the equilibrated solution was filtered through stacked 0.2 and 0.1µm Millex-VV Syringe Filter Units to remove cholesterol crystals.

### Polymer condensation in solution and at supported membranes

Phase boundaries were determined by titrating soluble polymers at constant NaCl concentration at constant temperature while monitoring the distribution of fluorescent signals by microscopy at 37°C. Polyelectrolyte concentration was titrated by adding small volumes of concentrated stocks to achieve the desired final concentration in solution. Systems exhibiting puncta in the majority of fields of view were assigned to contain coexisting phases. In some experiments, NaCl concentration was changed acutely by adding a small volume of a concentrated solution at the microscope stage. Cholesterol was added or removed from supported membranes by adding 150µl of either MβCD or MβCD-Chol with added polymers to 150µl of polymer solutions on the microscope stage. In absorption measurements, bright puncta visible in pGlu-532 images were segmented using a threshold whose level was set at 3 times the standard deviation of the first level wavelet of individual images. The total area fraction of segments (not shown) depended on details of how the binarization was accomplished, but the average intensity of segmented regions was robust to segmentation parameters.

### Preparation of U2OS cells

Human Bone Osteosarcoma Epithelial Cells (U2OS) (ATCC HTP-96) were obtained from Phyllis Hanson (University of Michigan, Ann Arbor MI). Cells were transiently transfected with plasmid-DNA encoding pCMV-CIB1-mRFP1-MP (addgene #58367)(32), pCMV-SNAP-CRY2-VHH(GFP) (addgene #58370)(35) and either PM-GFP (a gift from Barbara Baird and David Holowka, Cornell University, Ithaca, NY) (68), soluble eGFP (Lonza), pAc-GFPC1-Sec61beta (addgene #15108) (69) and/or CFP-Caax (addgene #15523) (70). Transfection was accomplished using Nucleofector electroporation (Lonza, Basel, Switzerland) with electroporation program CM-104. Generally, 500K cells were transfected with 1.5µg of total plasmid DNA and incubated overnight in glass-bottom culture dishes (P35G-1.5-10) (Mattek; Massachusetts, USA). For measurements involving multiple conditions that are compared directly, transfected cells are pooled prior to plating into dishes. Prior to an experiment, cells were labeled with the SNAP ligand, SNAP-Cell 647-SiR, following the manufacturer protocol. For fixed cell measurements, cells in imaging buffer (119 mM NaCl, 5 mM KCl, 25 mM HEPES buffer, 2 mM CaCl2, 2 mM MgCl2, 6 g/liter glucose, pH to 7.4) were exposed to controlled doses of blue light on a invitrogen gel imager (Safe Imager™ 2.0, Cat# G6600) prior to chemical fixation in 2% formaldehyde with 0.15% Glutaraldehyde for 10 min at room temperature. The fixative was then quenched with 3% bovine serum albumin (BSA) and imaged in the imaging buffer. In live cell experiments, cells in the imaging buffer were exposed to blue light through the objective using the epi-fluorescence light-source immediately prior to imaging.

For fixed cell measurements involving temperature perturbations, the light box was placed inside an incubator (37°C) or a cold room (4°C). Cells were equilibrated at the treatment temperature for 15 min prior to light exposure and fixation. For fixed cells pre-treated with n-alcohols, 5uL of concentrated DMSO stocks of either 1-octanol (120mM), 1-hexadecanol (6mM), or DMSO alone were diluted directly into 2ml of imaging buffer.

Cells were incubated at room temperature for 5 min before light exposure and chemical fixation. In the case of MβCD treatment, cells were incubated with 10 mM of MβCD at room temperature for 10 minutes prior to the light exposure and fixation. For MβCD/Chol treatment,saturated MβCD/Chol mixture was directly added to the cells without dilution and incubated at room temperature for 10 minutes followed by desired light exposure and fixation. For fixed cell measurements involving ionophore A23187 treatment, cells were incubated with 10μM of ionophore in the imaging buffer for 3 minutes prior to light exposure and chemical fixation. In separate measurements, PS exposure was monitored by Annexin V (Thermofisher, Cat#A23204).

In live cell measurements with acute treatments, cells were exposed to light prior to imaging and treatment. Perturbations were added to dishes after light exposure at room temperature as described for fixed cell measurements, with the exception of cholesterol perturbations in which the described solution was diluted 1:1. For temperature manipulation in live cell measurements, cells were plated on smart substrate and imaged as described for supported bilayer measurements with temperature control. Temperature was lowered below ambient temperature using Cold Gun System (Exair, Cat# 5215) mounted on the microscope.

For measurements with fatty acid feeding, stock solutions of palmitic acid (PA) or oleic acid (OA) in ethanol were mixed with 10% solutions of BSA in PBS at a 3:1 molar ratio then incubated at 37°C for 2h with continuous mixing (71). Solutions were diluted to 0.5mM fatty acid in culture media then sterile-filtered using a 0.22 µm syringe filter prior to incubation with cells for either 3h or 24h at 37°C. Control cells were incubated in media supplemented with BSA.

### Fluorescence imaging

Planar supported bilayers were imaged using an Olympus IX83-XDC inverted microscope with 100X UAPO TIRF objective (NA = 1.49), and active Z-drift correction (ZDC) (Olympus America). Fluorescence excitation was performed using LED lamp and fluorescence emission was detected on an EMCCD camera (Ultra 897, Andor). Typically, measurements were performed within a temperature controlled imaging chamber using a VAHEAT stage (Interherence) with PDMS (polydimethylsiloxane) reservoir held at 37°C. Different color channels were isolated using appropriate filter cubes (Chroma,Bellows Falls, VT USA). Single pLys-647 polymers were detected through total internal reflection illumination using a 647nm laser (OBIS 647, 120mW, Coherent, Santa Clara, CA) and single molecules were localized and tracked as described previously (37).

Confocal images of chemically fixed cells were acquired using ZEIS LSM 980 with Airyscan 2 Confocal microscope with plan-apochromat 63x/1.40 oil DIC M27 objective and GaAsP-PMT detector. Diffraction limited images of GPMVs and cells were acquired on the IX83 microscope described above or on a similarly equipped Olympus IX81 microscope.

### Quantifying cells containing puncta in fixed cell measurements

Multiple 82μm by 82μm fields of view (typically 20) within dishes were imaged in 3 colors to separately detect Cry2 (red excitation), CIB1-MP (green excitation), and GFP (blue excitation). Fields of view were collected randomly across the dish and typically contained 1-10 cells per field. After collecting an entire dataset spanning different treatments and light exposures, fields of view were randomized and cells within images were manually annotated by a separate individual blinded to the experimental conditions. Cells contained a broad distribution of expression levels and all cells expressing detectable levels of any labeled protein were annotated. Cells were determined to contain puncta if both the Cry2 and CIB1 channels exhibited multiple colocalized clusters. Error bounds (*σ*) for individual measurements arise from counting statistics and were tabulated as follows:

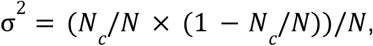

where N is the total number of cells annotated in the sample and Nc is the number of cells assigned as containing clusters. When multiple replicates are presented, points represent the weighted mean, with inverse variance weighting according to:

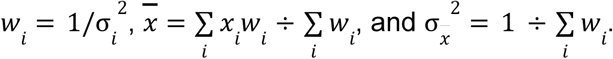

Points representing the fraction of cells containing clusters were fit to:

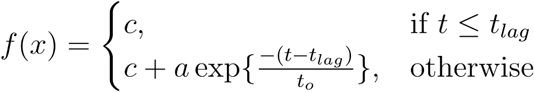

### Measuring Tmix in GPMVs

U2OS cells were labeled for 10 minutes with 2µg/mL DiI-C₁₂ then GPMVs were prepared using a vesiculation buffer containing 2 mM dithiothreitol (DTT) and 25 mM paraformaldehyde (PFA) as described previously. When specified, 300 μM 1-octanol or 15 μM 1-hexadecanol was added directly to prepared vesicles or included in the vesiculation buffer from stock solutions in DMSO. Transition temperatures were measured by imaging vesicles over a range of temperatures with a home-built temperature stage, tabulating the fraction of vesicles that are phase separated within images, then fitting to a sigmoid function to determine the temperature where 50% of GPMVs are phase separated.

### Dynamic light scattering (DLS)

Measurements were conducted using the DynaPro® NanoStar instrument (Wyatt Technology, Santa Barbara, CA, USA, Model WDPN-10, Serial# 410DPN) and analyzed using the DYNAMICS® software (Wyatt Technology). LUVs containing 0.02% DiO, 96% DOPC, 4% DOPE-DBCO or 0.02% DiO, 36% DOPC, 4% DOPE-DBCO, 40% DPPC, and 20% Chol were modified with pLys-N_3_. LUVs were placed in disposable cuvette without dilution with approximately 1.5mg/ml total lipids in phosphate buffer containing 400mM NaCl were placed in disposable cuvettes, and measurements were carried out at 37 ± 0.1°C. Soluble pLys and pGly polymers were titrated into the cuvette from concentrated stock solutions to achieve the concentrations specified in the text. At least three replicates per sample were performed, and for each replicate, 10 acquisitions of 5 seconds were collected to ensure consistency. The hydrodynamic diameter of the vesicles was determined using cumulant analysis within the instrument software. The average particle size was calculated as centroid of the diameter distribution as 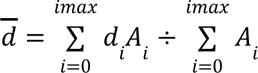 where *A_i_* is the amplitude of the histogram bin corresponding to diameter *d_i_*.

### Fluorescence Cross-Correlation Spectroscopy (FCCS)

Measurements were performed using ISS Alba time-resolved laser-scanning confocal system on IX-81 microscope (Olympus), with 60X water (UPLSAPO60XW), 1.2 NA objective in confocal volume equal to 1.26fl. Fluorescence excitation was performed using SC-400-6-PP supercontinuum laser (Fianium) with 6W integrated average power, ∼100ps pulse width, 20 MHz repetition rate and 400-2500nm wavelength range. A SPC-830 time-correlated single photon counting (TCSPC) board (Becker & Hickl) was used to collect photons. LUVs were prepared as in the DLS measurements, but pools of the same vesicle composition were labeled with different fluorophores (DiD, DiI, or DiO), and differently colored vesicles were pooled prior to conjugation with pLys-N_3_. Modified LUVs were diluted 1:100 (approx 15μg/ml) in a phosphate buffer with 400mM NaCl. LUV/polymer suspensions were contained within a temperature-controlled imaging chamber using a VAHEAT stage (Interherence) with PDMS (polydimethylsiloxane) reservoir held at 37°C. Soluble pLys and pGly polymers were titrated into the cuvette from concentrated stock solutions to achieve the concentrations specified in the text. Correlation curves were generated from photon counting traces using custom routines in Matlab that binned counts at 50kHz and divided traces into 1s intervals for correlation analysis. Prior to averaging, segments are filtered to remove those whose autocorrelation amplitudes are more than three scaled median absolute deviations away from the median in either color channel using the isoutlier() function in Matlab. The correlation coefficient was calculated as the cross-correlation amplitude divided by the square root of each of the autocorrelation amplitudes.

### Flow cytometry

Expression levels were characterized using an Attune NxT flow cytometer (Thermo Fisher Scientific) equipped with 405 nm, 488 nm, 561 nm, and 647 nm laser lines and appropriate filters. Cells were transfected and labeled with SiR-SNAP ligands as described for microscopy experiments. Single intact cells were gated based on forward-scatter and side-scatter measurements to exclude dead cells and debris. Data was collected using Attune NxT software, exported as .fcs files and raw acquired values were processed in MATLAB. In figures, points reflect all cells measured without additional gating. Some panels show filtered results and the methods used to gate are described in figure captions.

### Simulation Framework

We performed Monte-Carlo simulations on a 3-Dimensional DxLxL lattice, with L being the lateral to the membrane and D axial. We impose periodic boundary conditions on the two lateral *L* dimensions, and free boundary conditions at the *boundaries* of the axial dimension. **In all of our simulations D=32 and L=64**. Our bulk system consists of two soluble polymers whose interactions we described by a simple Hamiltonian:

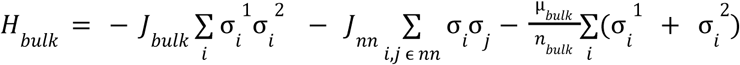

Here *J_bulk_* resembles the interaction strength between pLys and pGlu polymers in our experiments (tuned by salt concentration), J_m_ is a non-specific interaction, *μ_bulk_* is the chemical potential per bulk molecule, which sets the concentration, and *n_bulk_* = 10 is the number of monomers per bulk molecule. The superscripts 1 and 2 denote the two different polymer species that interact with each other, but self avoid.

We simulate a ‘grand-canonical ensemble’ in all simulations except the ‘Wetting’ simulation, where it is held constant. To keep the chemical potential in the bulk fixed, and sample many configurations of our bulk polymers, we separately simulate a ‘reservoir’ of bulk particles. At each monte-carlo step, we propose exchanges between the system and the reservoir. Particle swaps from the system to the reservoir simply delete the particle in the system. Particle swaps from the reservoir to the system duplicate the reservoir particle into the system.

We model a lipid membrane on the *z=0* boundary of our simulations. We add interactions with the membrane through a ‘tethered’ polymer component held at fixed concentration and restricted to diffusion in 2D in the membrane plane. These tethers resemble the 2^nd^ bulk polymer species above, except the length of tethers is n_tether_ = 5 monomers. The interactions between the tethered polymers and the bulk are captured by the Hamiltonian:

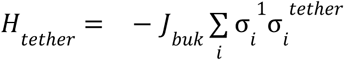

In simulations where we model a multicomponent membrane, we additionally include an ising model at the z*=0* boundary of the simulation box. The energetics of the Ising model are described by the Hamiltonian:

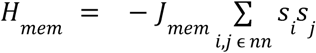

where s_i_ and s_j_ are spins at lattice positions i and j, and the sum runs over all nearest-neighbors. Here *J_mem_* sets the strength of interactions in the Ising model, and we fix the magnetization and thus have no magnetic field term. We simulate both tethers and ising spins at fixed composition.

### Simulation Procedure

We sample configurations of these simulations with a Monte-Carlo procedure. Bulk molecules are free to move in the simulation box, and their configurations are updated with local ‘kink’ and ‘reptation’ moves. These moves are accepted with the Metropolis probability *p_accept_ = max(e^βH^,1)*. We also propose translations of clusters of bulk molecules. We only accept these cluster moves if no bonds are formed or broken during the move, following detailed balance. We exchange bulk molecules with a particle reservoir, whose acceptance probability is set by the Hamiltonian above.

We update tether molecules with translations in the plane of the membrane, which are again accepted with a Metropolis probability. In simulations modeling membranes with immobile tethers we do not propose any tether moves. In simulations with the Ising membrane, the tether can only move on s_i_= -1 membrane spins, to mimic the coupling to a specific membrane phase. Likewise, we reject any ising spin-exchanges that attempt to flip a spin on the same position as a tether. We simulate the ising model with conserved-spin, local Kawasaki dynamics where spin-exchanges are only proposed among nearest-neighbors. Each Monte-Carlo step of our simulation proposes updates for every particle and spin, and a number of particle exchanges with the reservoir. We ran simulations up to ∼10^8^ Monte-Carlo steps to ensure equilibration.

Adsorption Isotherms To calculate the adsorption in our simulations, we first identify whether a prewet phase is present, and then calculate adsorption in each region (prewet or dry) separately. To determine whether or not the system is prewet, we time-averaged 10 simulation configurations over a window of 5x10^5^ monte-carlo steps, and smooth simulations over a 4x4x1 window. Regions where the time-averaged density of tether and bulk species exceed 1.0 are declared ‘prewet’, and all other regions to be ‘Dry’. We declared simulations where the average fraction of prewet domain exceeded 16/64^2^, the length scale set by our smoothing step, to be ‘Prewet’. We then identify *dense-phase* domains as lattice points where the time-averaged density of tether and bulk molecules is greater than or equal to a threshold value of 1. After identifying these regions we average the density in the dense phase over the two lateral dimensions at each *z* coordinate, and then sum the final density over *z.* When the average area fraction of domains is below 16/64^2^ we declare the system as *single-phase*, and average the density over all lateral coordinates before summing over *z.* For immobile tether simulations, we average these results over ten random tether configurations.

## ACKNOWLEDGEMENTS

We thank Guoming Gao for assistance with some early experiments, and Andrea Stoddard for support with molecular biology and cell culture. Research was supported through grants from the NSF (1808551 to BBM and SLV) and the NIH (GM138341 to BBM and GM152150 to SLV).

## DATA AVAILABILITY STATEMENT

Raw data, including microscopy images, will be made available on public servers when published.

