## Supplemental movies, figures, and tables for "The membrane transition strongly enhances biopolymer condensation through prewetting"

### SUPPLEMENTARY MOVIE CAPTIONS

Supplementary Movie 1: **Diffusion and coarsening of prewet polyelectrolyte domains.** Membranes contain 96% DOPC and 4% DOPE-pLys and the solution contains 400mM NaCl and 5μg/ml pLys/pGlu at 1:1 w/w. Domains move and merge on defect-free regions of the membrane. Supported membranes prepared through spin-coating and hydration often contain linear (one dimensional, 1D) defects. Prewet domains tend to wet these 1D defects, and merging events of prewet domains absorbed to 1D defects can be seen in this video. Scale bar is 10μm in the main frame and 2μm in the inset.

Supplementary Movie 2: **Domain melting with added NaCl.** Membranes contain 96% DOPC and 4% DOPE-pLys and the solution contains 10μg/ml pLys/pGlu at 1:1 w/w. Initially, NaCl concentration is 150mM, and domains melt after the addition of NaCl to make a 2M solution. Scale bar is 10μm in the main frame and 2μm in the inset.

Supplementary Movie 3: **Domain melting by removing soluble polyelectrolytes from solution.** Membranes contain 96% DOPC and 4% DOPE-pLys. The solution initially contains 400mM NaCl with 5μg/ml pLys/pGlu at 1:1 w/w, then is replaced with a solution of 400mM NaCl without polymers. Scale bar is 10μm in the main frame and 2μm in the inset.

Supplementary Movie 4**: Reversible membrane immiscibility transition in a planar bilayer stack.** A supported bilayer stack composed of 36% DOPC, 40% DPPC, 20% Chol, 4% DOPE-DBCO, and 0.2% DiO was prepared via spin coating of lipids from solvent followed by hydration in 150mM PBS. Initially the bilayer is in a single liquid phase at 37°C, then phase separation occurs when temperature is lowered to 32°C. Temperature is cycled twice between these values over the time-course.

Supplementary Movie 5: **Domain melting with cholesterol addition.** Membranes of 36% DOPC, 40% DPPC, 20% Chol, and 4% DOPE-pLys in the presence of 0.25μg/ml pLys/pGlu and 400mM NaCl contain prewet domains. Domains melt after 2.5mg/ml MβCD-Chol is added to the solution while maintaining the concentration of soluble polymers.

Supplementary Movie 6: **Removing Chol from a 40% Chol membrane.** Membranes of 16% DOPC, 20% DPPC, 40% Chol, and 4% DOPE-pLys in the presence of 0.5μg/ml pLys/pGlu and 400mM NaCl do not contain prewet domains. Domains form after 10mM MβCD is added to the solution while maintaining the concentration of soluble polymers.

Supplementary Movie 7**: Diffusion and coarsening of prewet protein domains at the plasma membrane in U2OS cells.** A U2OS cell expressing Cry2-αGFP, CIB1-MP, and PM-GFP. Cry2-αGFP is labeled with the SiR-SNAP tag ligand which is visualized immediately after a transient exposure to blue light at 37°C. Some puncta are out of focus because the ventral surface is imaged. Domains are circular, highly mobile, and coarsen via coalescence. Scale bar is 10μm in the main frame and 2μm in the inset.

Supplementary Movie 8**: Prewet domains dissipating over time in the absence of blue light illumination.** A U2OS cell expressing Cry2-αGFP, CIB1-MP, and PM-GFP. Cry2-αGFP is labeled with the SiR-SNAP tag ligand which is visualized immediately after a transient exposure to blue light at 22°C. The majority of domains dissociate over a span of 8 minutes.

Supplementary Movie 9: **Prewet domains in U2OS cells form upon temperature quench.** A U2OS cell expressing Cry2-αGFP, CIB1-MP, and PM-GFP. Cry2-αGFP is labeled with the SiR-SNAP tag ligand which is visualized immediately after a transient exposure to blue light at 37°C. Initially, Cry2-αGFP is uniformly distributed, but condenses into mobile puncta after temperature is quenched to 15°C. Scale bar is 10μm in the main frame and 2μm in the inset.

Supplementary Movie 10: **Prewet domains in U2OS cells melt upon treatment with 300mM octanol.** A U2OS cell expressing Cry2-αGFP, CIB1-MP, and PM-GFP. Cry2-αGFP is labeled with the SiR-SNAP tag ligand which is visualized immediately after a transient exposure to blue light at 22°C. When indicated, 300mM octanol is added to the solution. Puncta dissipate and the overall brightness of Cry2-αGFP at the membrane diminishes. Scale bar is 10μm in the main frame and 2μm in the inset.

Supplementary Movie 11: **Prewet domains in U2OS cells form upon treatment with M□CD-Chol.** A U2OS cell expressing Cry2-αGFP, CIB1-MP, and PM-GFP. Cry2-αGFP is labeled with the SiR-SNAP tag ligand which is visualized immediately after a transient exposure to blue light at 22°C. When indicated, 2.5mg/ml M□CD-Chol is added to the solution. Initially Cry2-αGFP is uniformly distributed, then mobile Cry2-αGFP puncta assemble upon M□CD-Chol addition. Scale bar is 10μm in the main frame and 2μm in the inset.

Supplementary Movie 12: **Prewet domains in U2OS cells melt upon treatment with M□CD.** A U2OS cell expressing Cry2-αGFP, CIB1-MP, and PM-GFP. Cry2-αGFP is labeled with the SiR-SNAP tag ligand which is visualized immediately after a transient exposure to blue light at 22°C. When indicated, 5mg/ml M□CD is added to the solution. Puncta dissipate and the overall brightness of Cry2-αGFP at the membrane diminishes. Scale bar is 10μm in the main frame and 2μm in the inset.

Supplementary Movie 13: **Prewet domains in U2OS cells melt upon treatment with A23187.** A U2OS cell expressing Cry2-αGFP, CIB1-MP, and PM-GFP. Cry2-αGFP is labeled with the SiR-SNAP tag ligand which is visualized immediately after a transient exposure to blue light at 22°C. When indicated, 10μM A23187 is added to the solution. Puncta dissipate and the overall brightness of Cry2-αGFP at the membrane diminishes. Scale bar is 10μm in the main frame and 2μm in the inset.

Supplementary Movie 14: **Diffusion and coarsening of prewet protein domains at the ER membrane in U2OS cells.** A U2OS cell expressing Cry2-αGFP, CIB1-MP, and Sec61-GFP. Cry2-αGFP is labeled with the SiR-SNAP tag ligand which is visualized immediately after a transient exposure to blue light at 37°C. Scale bar is 10μm in the main frame and 2μm in the inset.

Supplementary Movie 15: **Prewet domains at the ER in U2OS cells form upon treatment with M□CD-Chol.** A U2OS cell expressing Cry2-αGFP, CIB1-MP, and Sec61-GFP. Cry2-αGFP is labeled with the SiR-SNAP tag ligand which is visualized immediately after a transient exposure to blue light at 22°C. When indicated, 2.5mg/ml M□CD-Chol is added to the solution. Initially Cry2-αGFP is uniformly distributed, then mobile Cry2-αGFP puncta assemble upon M□CD-Chol addition. Scale bar is 10μm in the main frame and 2μm in the inset.

**SUPPLEMENTARY FIGURES AND CAPTIONS**

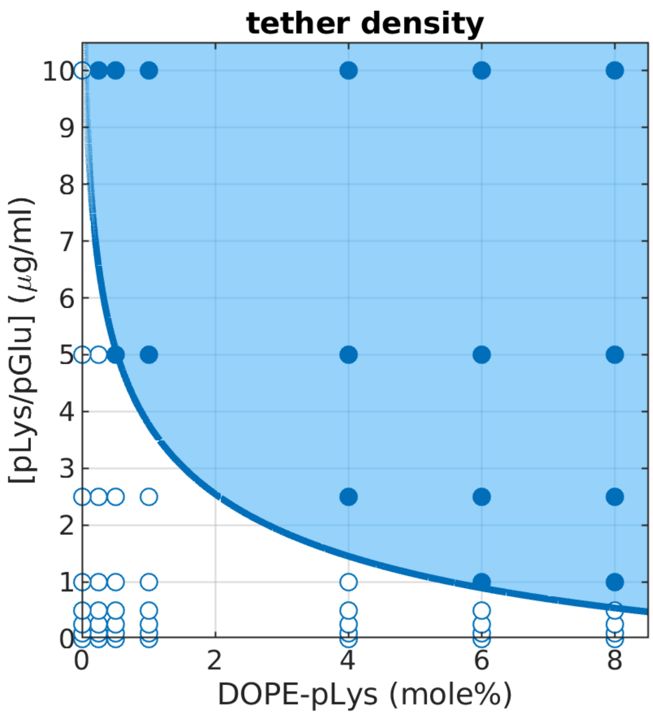


**Supplementary Figure S1:** Conditions where prewetting is detected on majority DOPC membranes with varying incorporation of DOPE-pLys at constant NaCl concentration (150mM). Open points indicate the presence of a single surface phase while closed points indicate the presence of 2 coexisting surface phases, one a prewet phase rich in soluble polymer and DOPE-pLys tethers. The blue line is drawn to guide the eye and is not a fit to any theory.

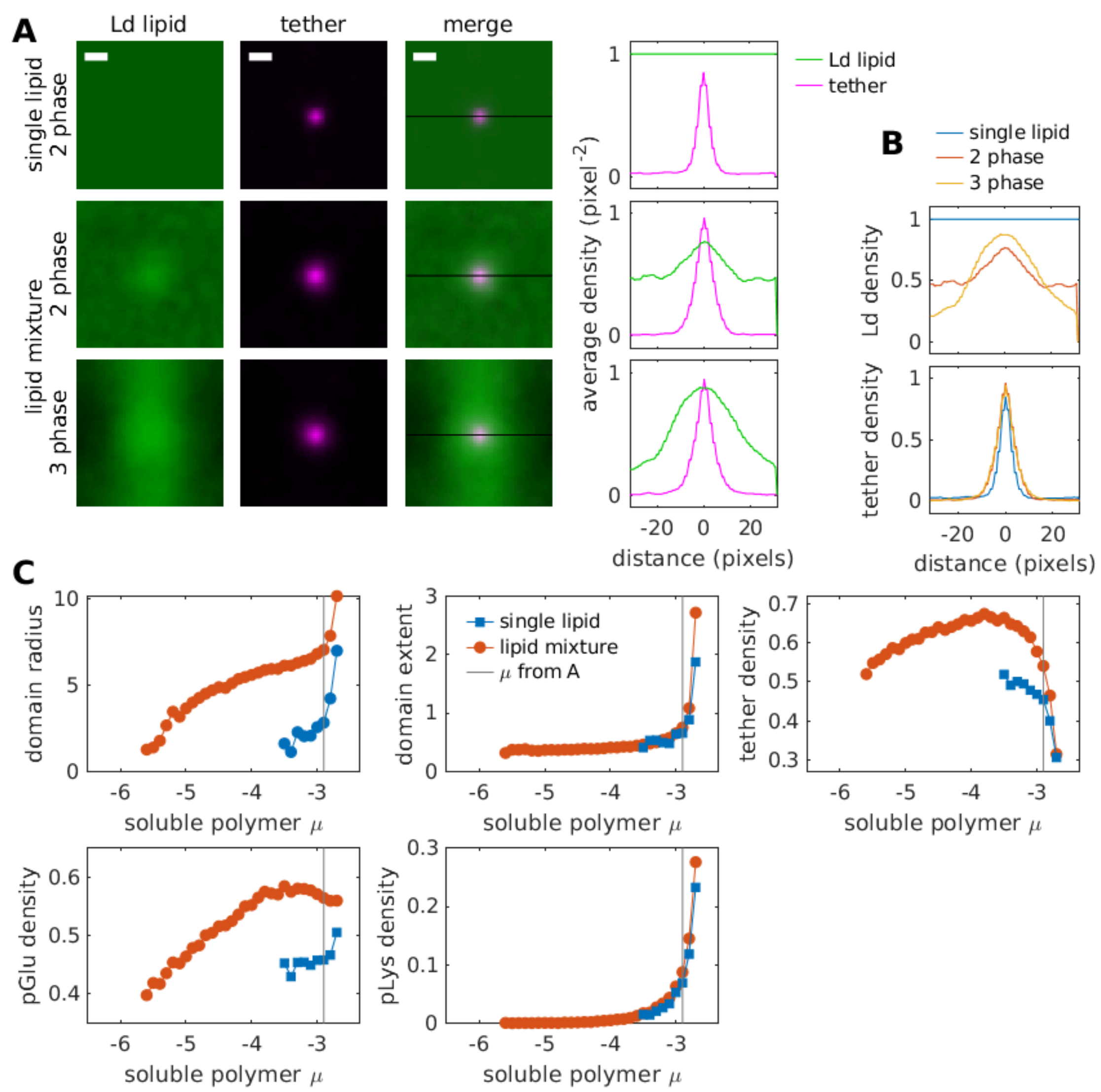


**Supplementary Figure S2:Properties of simulated prewet domains on single-component and multi-component membranes.** (A) Time-averaged configurations of membrane lipids (green) and tethers (magenta) for μ=-2.9. (right) Densities across a line-scan of the surface. (top) Surfaces containing a single lipid and tethers display 2 surface phases at prewetting, one phase rich in tethers and another phase depleted in tethers. Surfaces with multiple lipids can phase separate and display two- or three- phase coexistence. Prewet phases in the two-phase coexistence region are enriched in tethers and Ld lipids and the dry phase is depleted in tethers. In the three phase coexistence region, the prewet phase is enriched in tethers and Ld lipids. In addition there are 2 dry phases, one enriched in Ld lipids and one depleted in Ld lipids. (B) Comparison of lipid and tether densities across conditions replotted from part A: tethers are modestly more enriched in complex membranes. ( C) Properties of prewet domains as a function of bulk chemical potential. Domain radius, the two-dimensional size of prewet domains, is consistently larger on multicomponent membranes while the domain extent, the distance the droplet extends into the bulk, is nearly identical. This is consistent with the theoretical prediction that domain extent is dominated by bulk properties, and not surface properties. The tether density is consistently higher in multi-component membranes, and both membranes see a decrease in tether density approaching wetting. pGlu density is highly enriched at low chemical potentials in both membranes, while pLys density only increases near wetting.

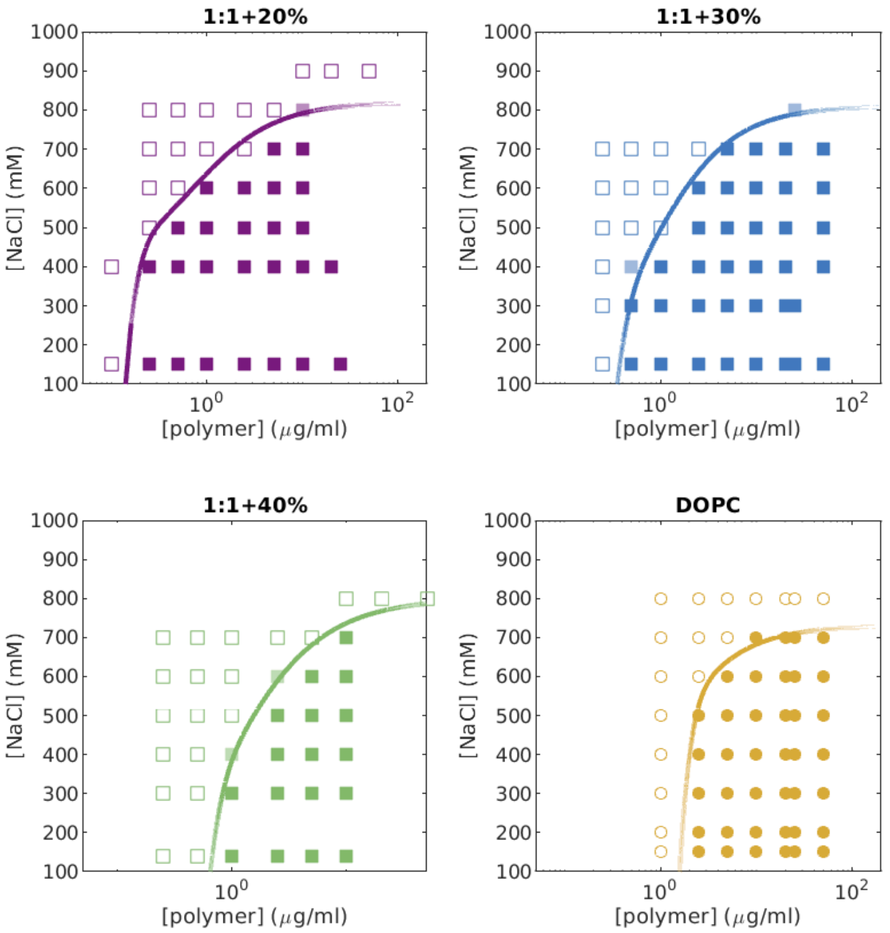


**Supplementary Figure S3:** Compositions and NaCl Concentrations imaged to obtain the phase diagram of Figure 2G. Open points indicate the presence of a single surface phase while closed points indicate the presence of 2 coexisting surface phases. The line is drawn to delineate the boundary between phase separated and uniform conditions and is not a fit to any model.

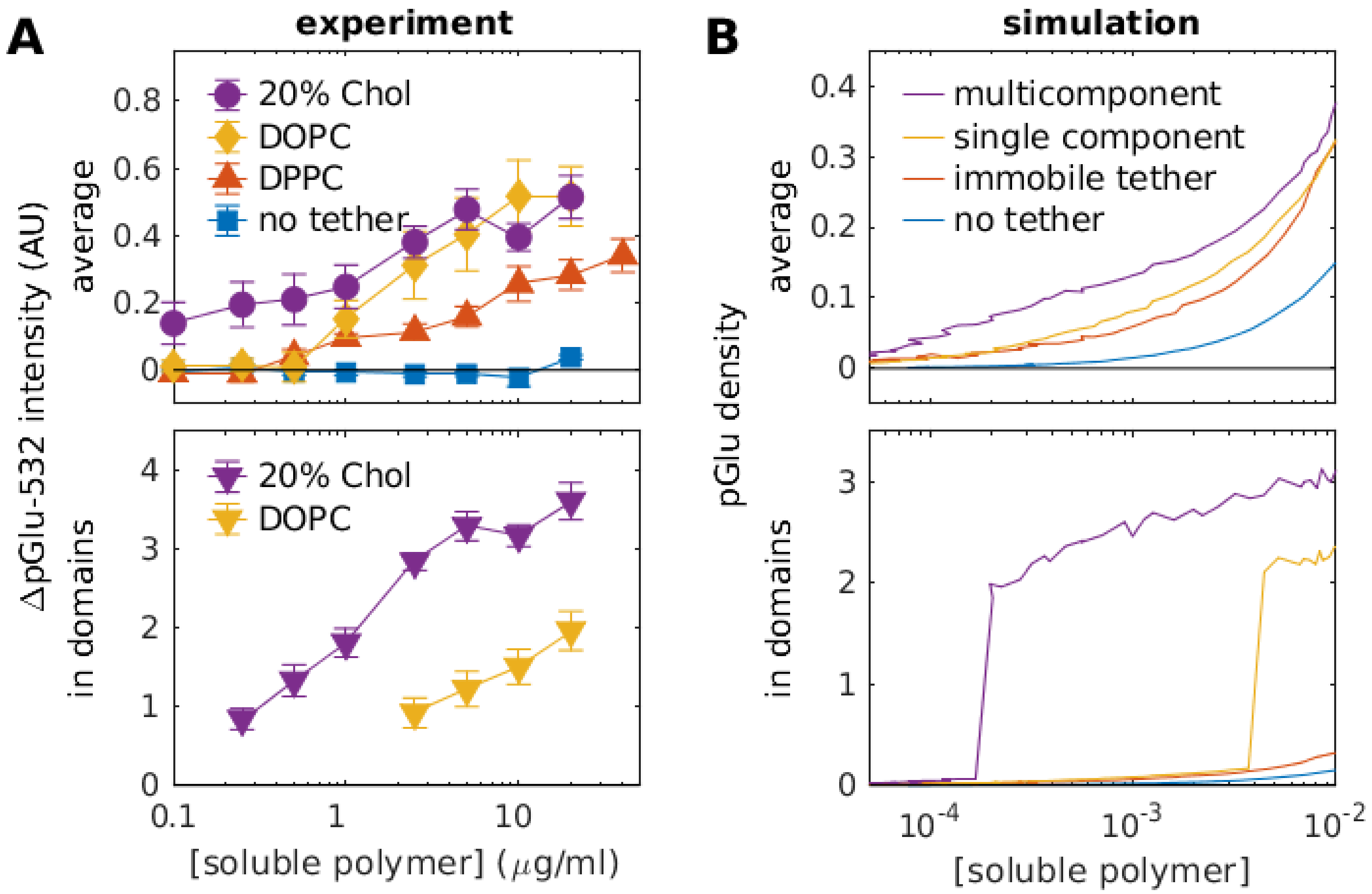


**Supplementary Figure S4: Prewetting and adsorption of polyelectrolytes on membranes in experiment and simulation.** (A) pGlu-532 fluorescence intensity observed in experiments. (Top) Average fluorescence intensity across multiple fields of view focused for each condition, with the objective focused on the membrane plane. (bottom) Average fluorescence intensity of prewet domains segmented from the full field of view as described in Methods. For all traces, the fluorescence intensity detected prior to adding soluble polymers was subtracted from each value. (B) Polymer concentrations extracted from Monte-Carlo simulations. (top) Average integrated concentration across the entire surface. (bottom) concentration in prewet domains only. Note that in simulations and experiments, only fluid, coupled surfaces (20% chol and DOPC) have prewet domains.

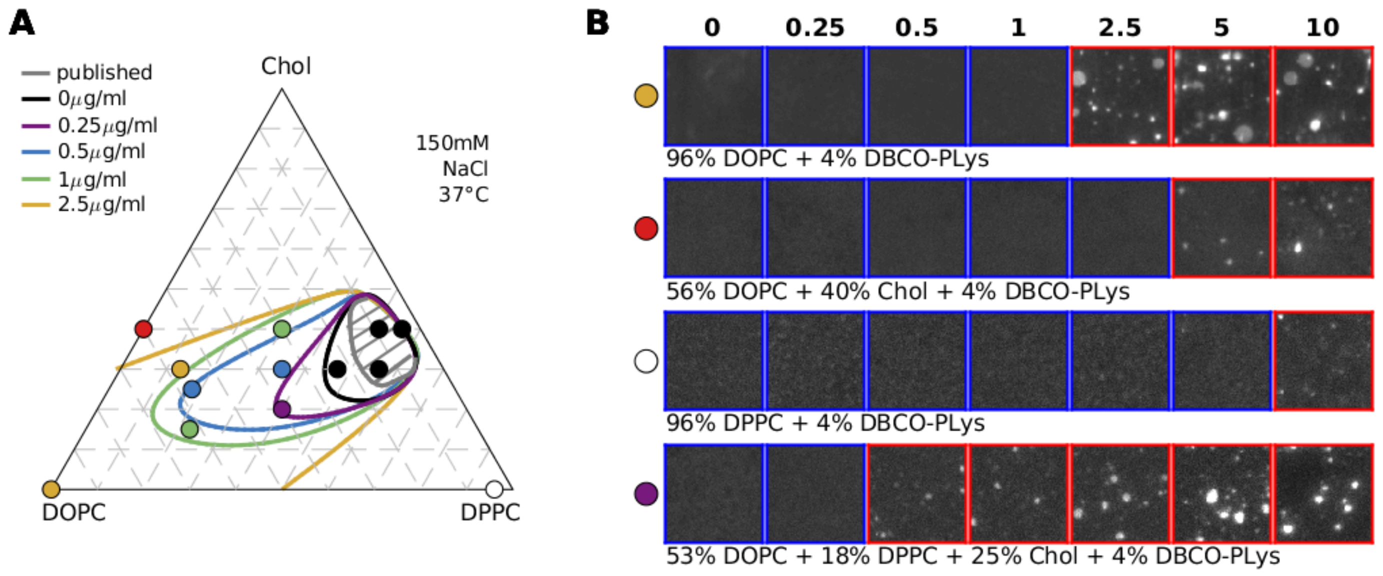


**Supplementary Figure S5:** Phase diagram and micrographs incorporating other membrane lipid compositions. (A) Phase diagram repeated from Fig 2H showing the composition of points interrogated to estimate the phase boundaries drawn. (B) Micrographs showing polyelectrolyte titrations on bilayers of the specified compositions used to determine $c_{prewet}$. While most membrane compositions interrogated potentiated prewetting as compared to the majority DOPC membrane, there were a few compositions where prewetting was inhibited. Membranes made of DPPC and DOPE-pLys are in a gel phase at 37°C, and the reduced mobility DOPE-pLys tethers acts to increase $c_{prewet}$ close to $c_{sat}$, as reported previously (17). $c_{prewet}$ is also elevated in membranes of DOPC, Chol, and DOPE-pLys compared to DOPC/DOPE-pLys membranes. Past work suggests that interactions between DOPC and Chol lipids are slightly less favorable than those between Chol-Chol interactions, which would lead to dispersion of DOPC lipids in these membranes. It is possible that prewetting is moderately inhibited in these membranes because this type of non-ideal mixing also gives rise to a membrane-mediated repulsion between DOPE-pLys tethers.

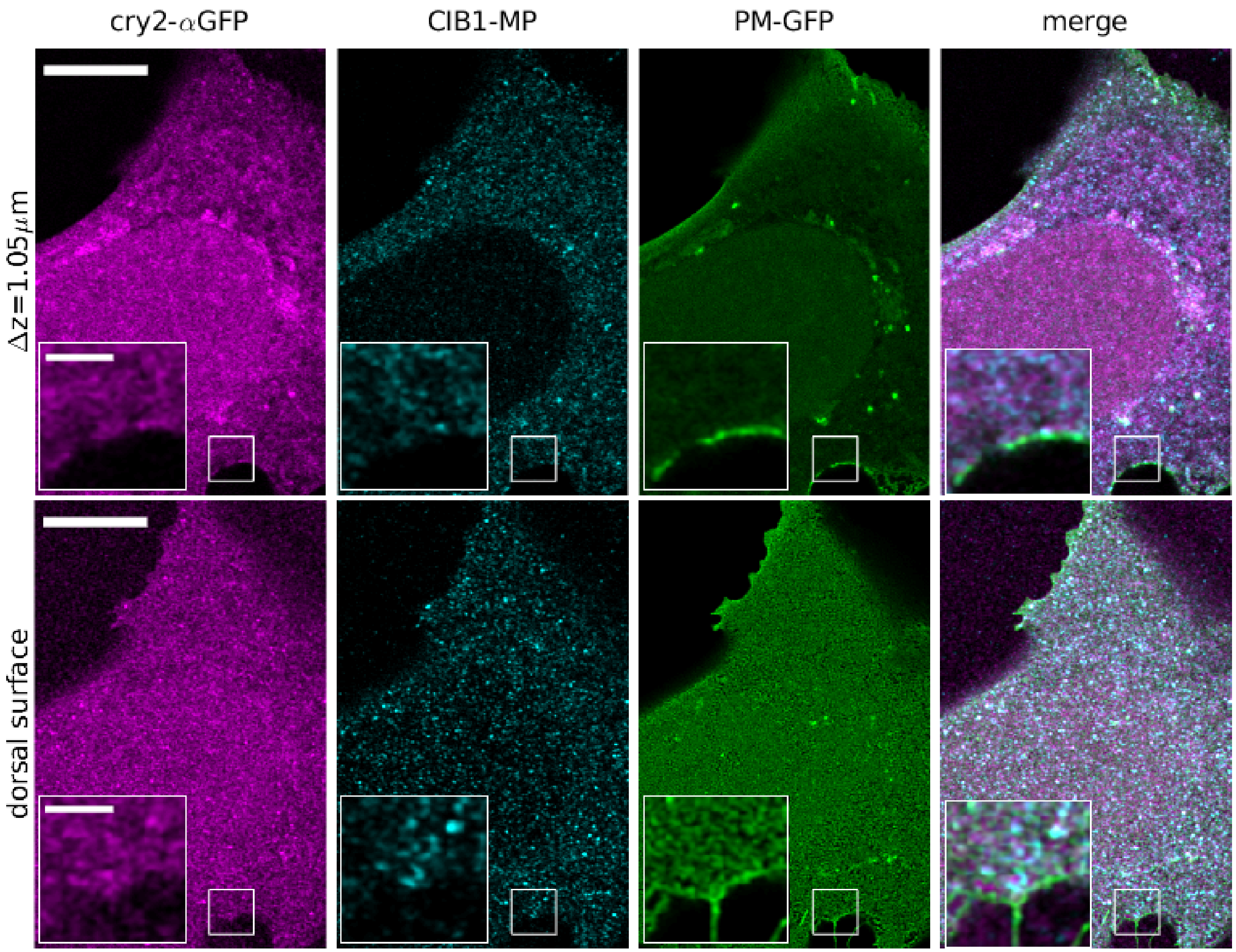


**Supplementary Figure S6:** Localization of expressed proteins prior to blue light exposure obtained by confocal imaging in representative cell.

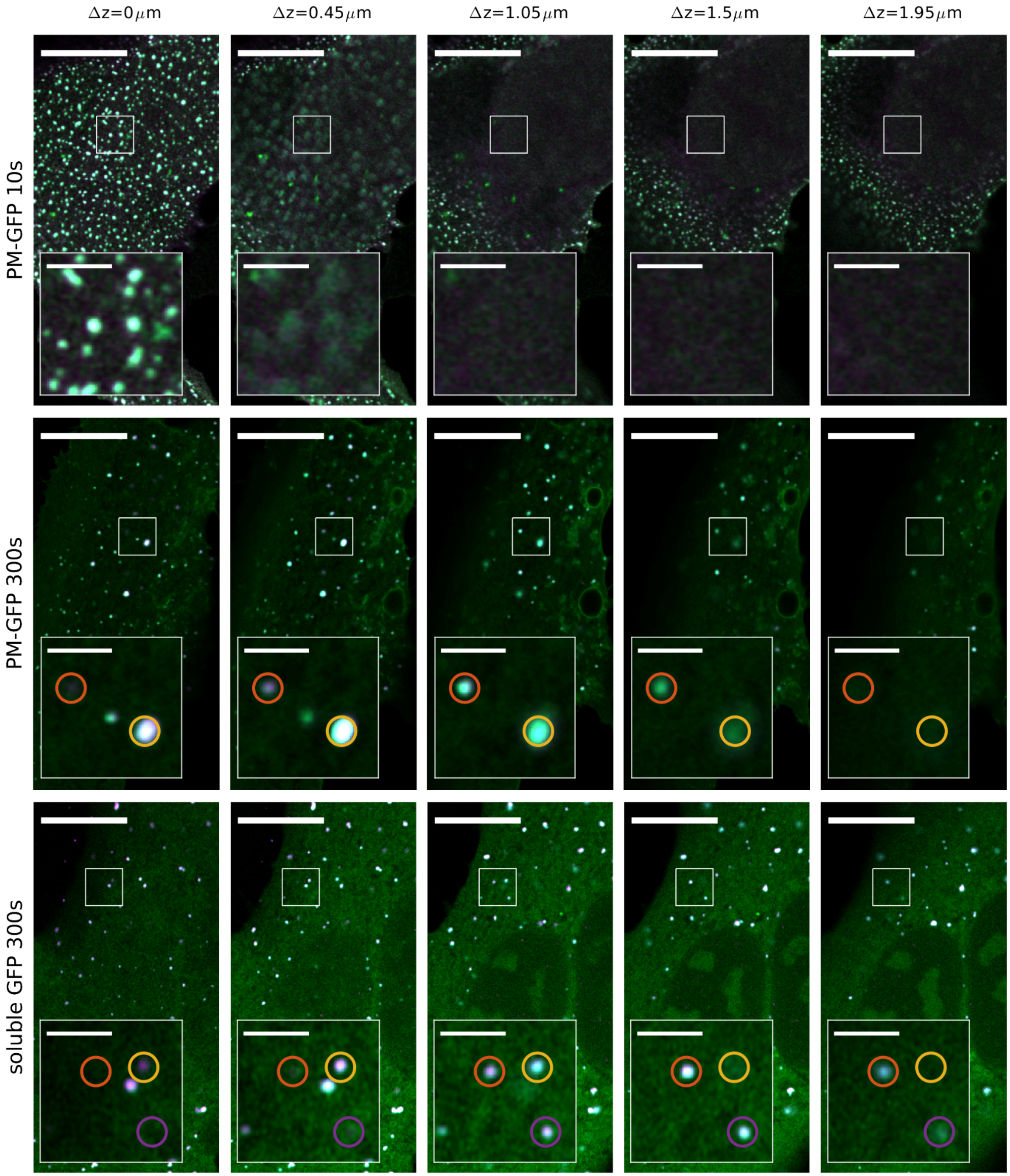


**Supplementary Figure S7: Examples of prewet (top), wet (middle), and dry (bottom) domains in confocal images of U2OS cells.** (Top) Upon short light exposures, plasma membrane localized puncta are not visible in confocal slices translated axially above the dorsal surface. (Middle) With long light exposures, plasma membrane puncta can extend into planes >1μm away from the dorsal surface. (top) Puncta formed with long light exposure with soluble GFP are present throughout the cytoplasm and the same puncta can appear in multiple axial planes.

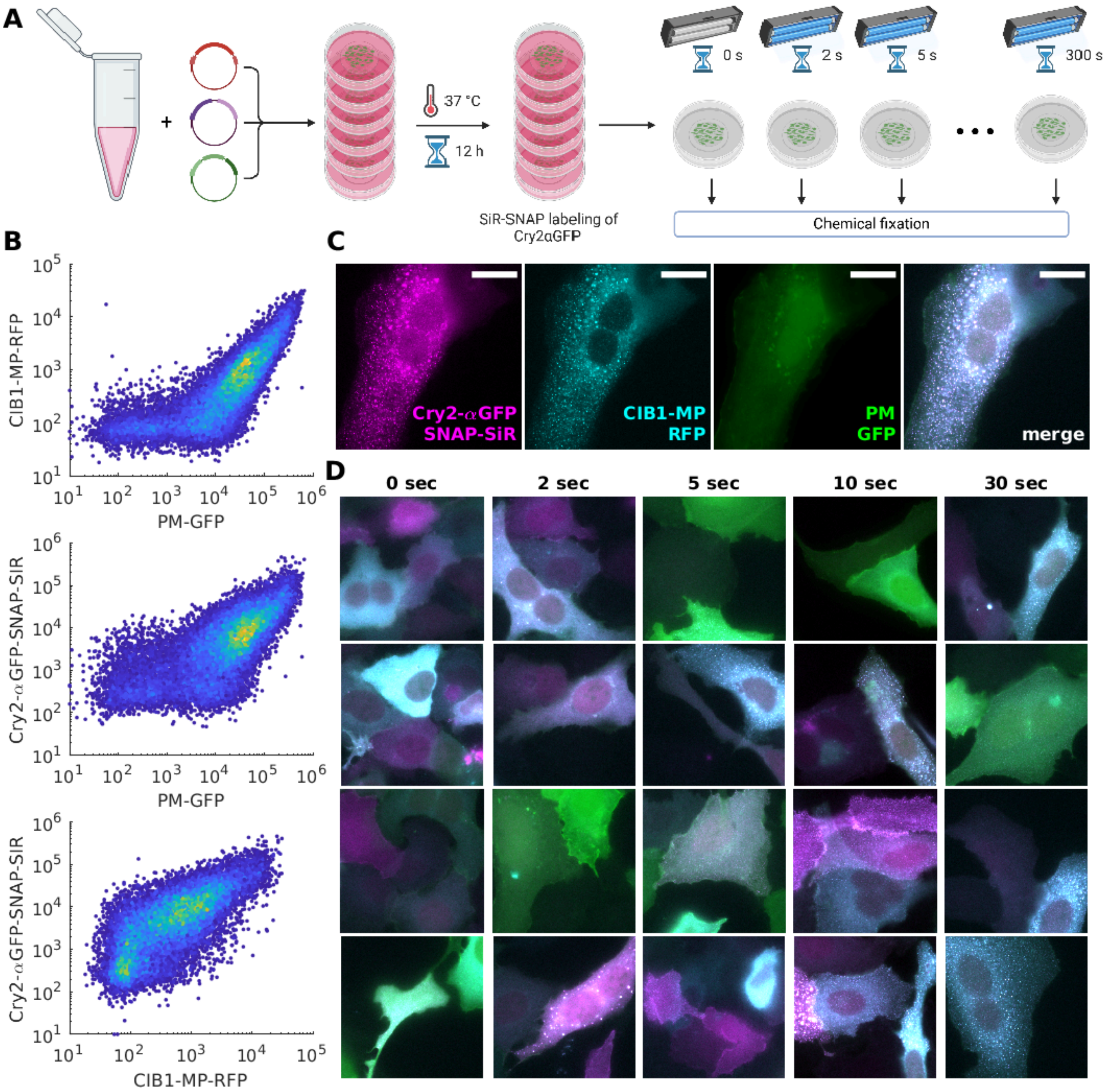


**Supplementary Figure S8: Overview of quantitative assay.** (A) Schematic representation of sample preparation workflow. U2OS cells electroporated with plasmids encoding Cry2-αGFP-SNAP, CIB1-MP-RFP, and PM-GFP were plated on Mattek dishes and incubated overnight prior to labeling with SiR-SNAP-ligands. Cells in individual dishes were then exposed to different durations of blue light then immediately chemically fixed prior to imaging. (B) Flow cytometry analysis of U2OS cells transfected under identical conditions show a broad range of expression levels of individual constructs, and that expression levels strongly correlate across expressed proteins.(C) A representative field of view showing individual color channels from a dish exposed to 30s of blue light at 22°C. This cell exhibits colocalized puncta in the Cry2-αGFP and CIB1-MP channels and is considered clustered even though puncta are not clearly visible in the PM-GFP channel. (D) Representative fields of view merging the three color channels for a range of exposure times from a single experiment. Typically at 20-30 fields of view are randomly acquired across a single dish for a single measurement and all cells expressing any fluorophore in the field of view are scored as being either clustered or nonclustered.

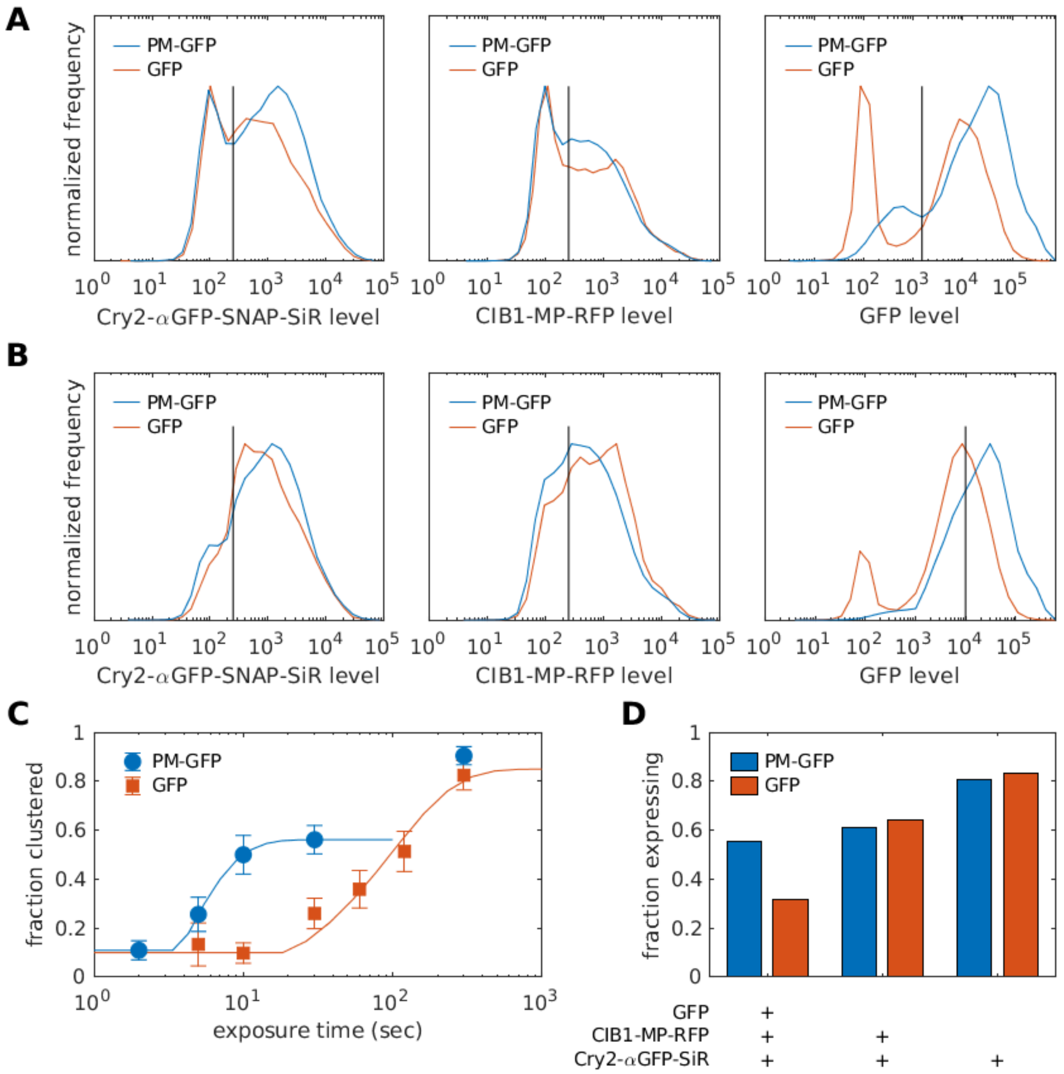


**Supplementary Figure S9: Expression of Cry2-αGFP-SNAP and CIB1-MP-RFP in U2OS cells also transfected with PM-GFP or GFP constructs.** (A) Comparison of fluorescence intensities of cells transfected with plasmids for Cry2-αGFP-SNAP, CIB1-MP-RFP, and either PM-GFP or soluble GFP. Cells are labeled with SiR-SNAP ligands prior to measurement. (B) Expression level distributions from A after filtering out cells with intensities below the threshold values indicated in Part A for all three channels, replicating the criteria for a cell to be considered in the microscopy assay described in Supplementary Figure S8. (C) Microscopy assay results for a titration of blue light exposures from cells taken from the same transfection as used to produce the distributions in A,B. Even though expression of Cry2-αGFP and CIB1-MP-RFP are similar across conditions, significantly more light is required to detect puncta in GFP expressing cells the microscopy assay. Error bars represent errors associated with counting statistics within this single replicate, tabulated as described in Methods. (D) Fractions of cells with fluorescent intensities above the threshold values indicated in part B for the indicated protein combinations. Values are normalized by the total cell population represented in part B.

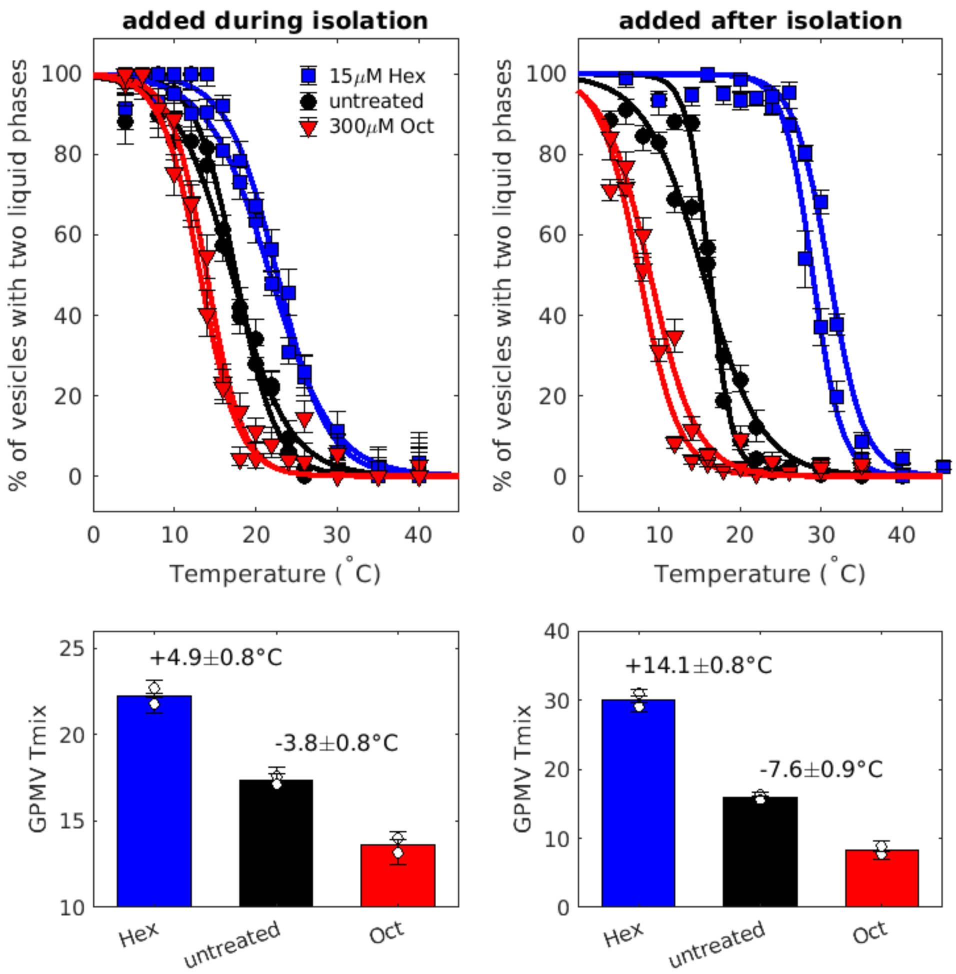


**Supplementary Figure S10: Giant plasma membrane vesicles (GPMVs) isolated from U2OS cells exhibit a miscibility transition that is modulated by octanol and hexadecanol.** (Top) Points showing the fraction of GPMVs that contain coexisting liquid phases as a function of temperature. Points are fit to sigmoid functions to determine the midpoint of the transition which is reported as Tmix (bottom). Treatment with hexadecanol and octanol either during GPMV isolation (left) or after isolation (right) raises and lowers GPMV Tmix respectively although with different magnitudes, likely due to the different lipid:alcohol ratios in the two treatment protocols. Error bars on individual points are tabulated from counting statistics. (bottom) The change in average Tmix for the treatments shown above, extracted by fitting measured curves to a sigmoid function. Error bars represent the SEM between replicates.

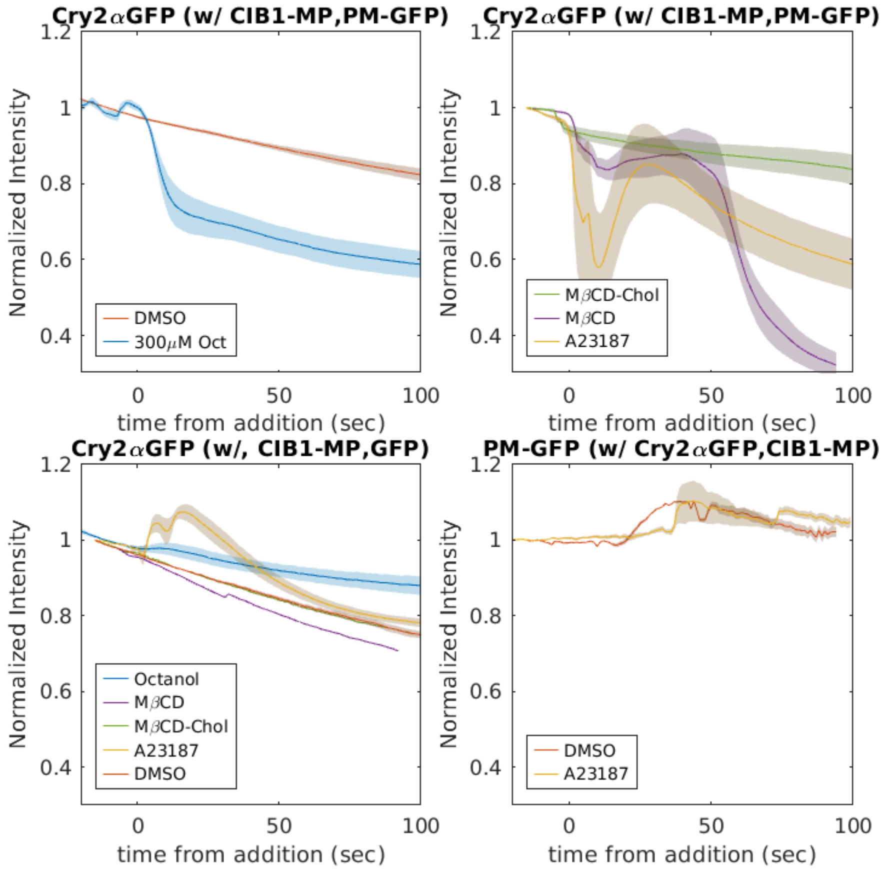


**Supplementary Figure S11:** Cry2-αGFP levels within the focal plane of the ventral plasma membrane are impacted by treatments in live U2OS cells. (Top left) Treatment with 300μM octanol leads to Cry2-αGFP desorption from the dorsal plasma membrane. (Top right) More complex desorption kinetics are observed for both MβCD and A23187 treatment, but levels remain largely unchanged with cholesterol loading with MβCD-Chol. (Bottom Left) Cry2-αGFP levels are largely unchanged by chemical treatments when co-expressed with soluble GFP and CIB1-MP. Probe photobleaching varies across treatments due to variation in expression levels. (Bottom right) PM-GFP levels are not reduced with A23187 treatment.

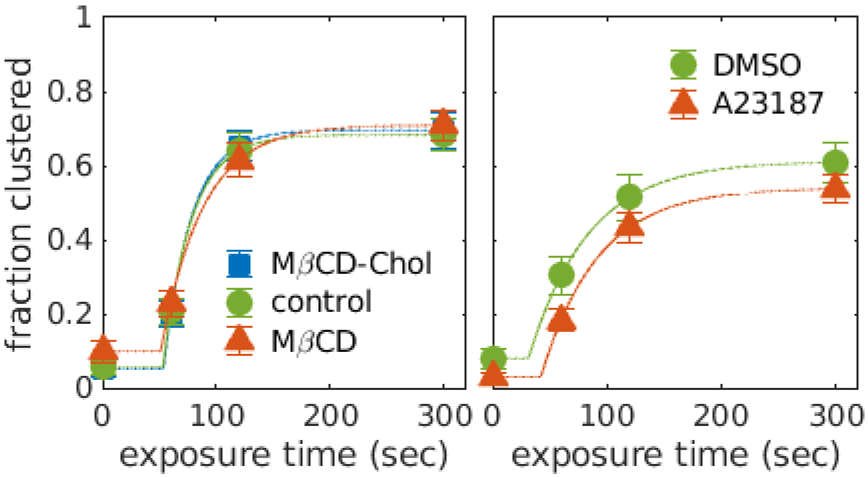


**Supplementary Figure S12: Cholesterol and A23187 treatment do not impact the stability of Cry2-CIB1 puncta assembled in the presence of soluble GFP.** Curves representing the percentages of cells exhibiting Cry2-αGFP/CIB1-MP puncta in fields of cells chemically fixed after the varying exposure times to blue light. Cells transiently express Cry2-αGFP, CIB1-MP, and a soluble form of GFP. (Left) Cells are pretreated for 5 min with either 10mg/ml M□CD or the M□CD-Chol solution described in Methods prior to light exposure and chemical fixation. (right) Cells were pretreated for 5 min with 10μM A23187 prior to light exposure and chemical fixation. Points represent a variance weighted mean over 3 replicates and error bars are the SE of the weighted mean tabulated as described in Methods.

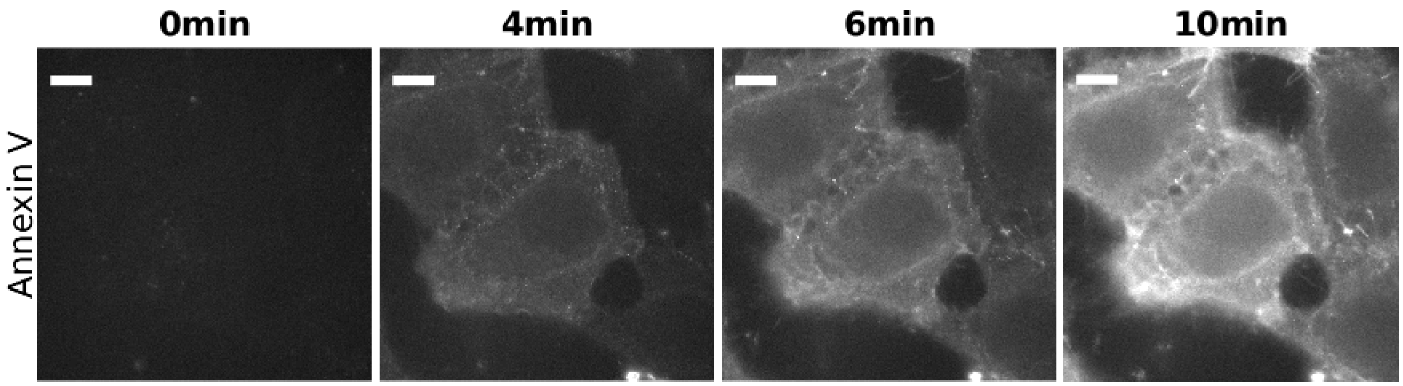


**Supplementary Figure S13. Cellular Anexin V staining of U2OS cells after treatment with 10μM of A23187.**
Scale-bar is 10μm.

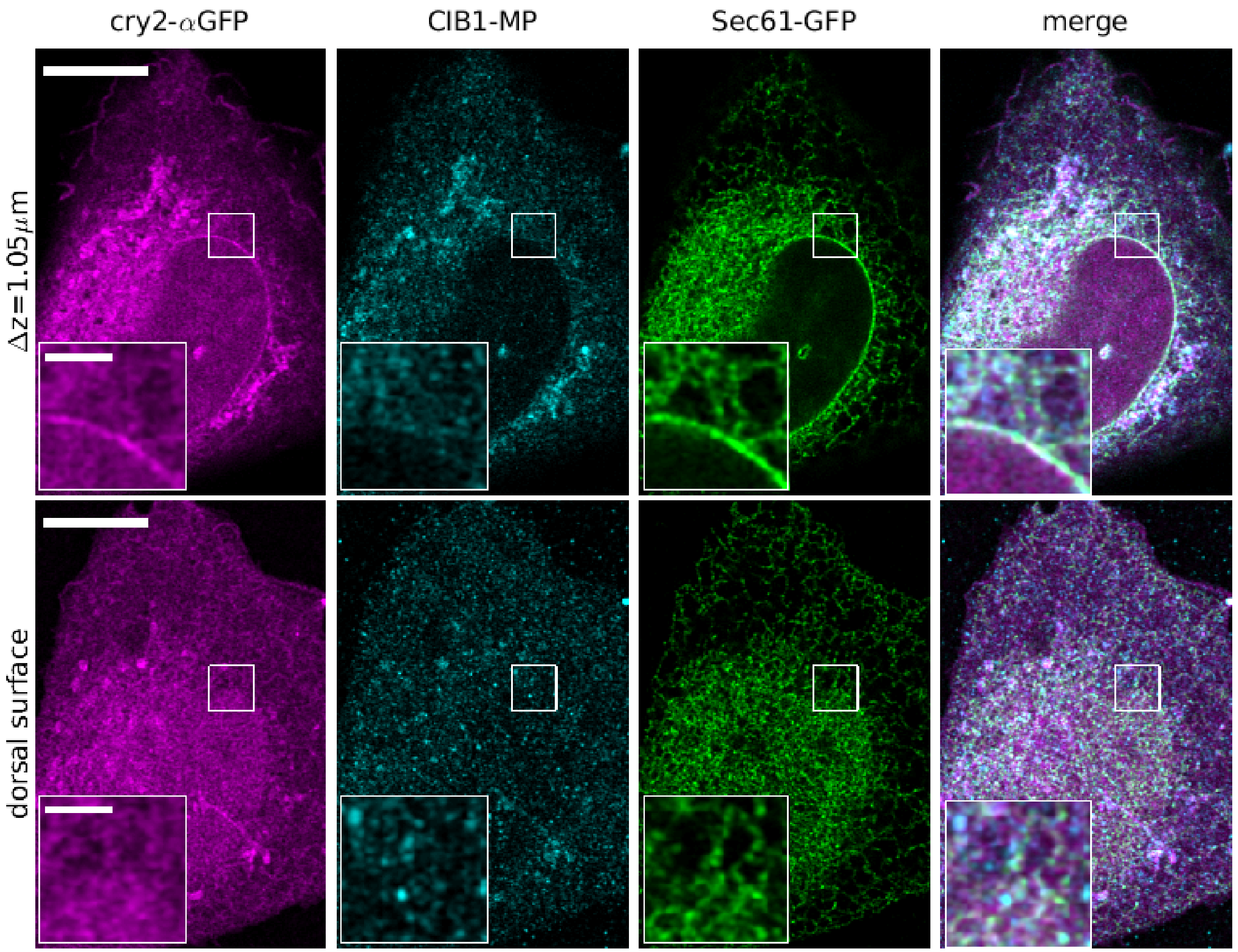


**Supplementary Figure S14: Localization of expressed proteins prior to blue light exposure obtained by Airyscan confocal imaging in representative U2OS cell transiently expressing Cry2-αGFP and CIB1-MP and Sec61 GFP.**

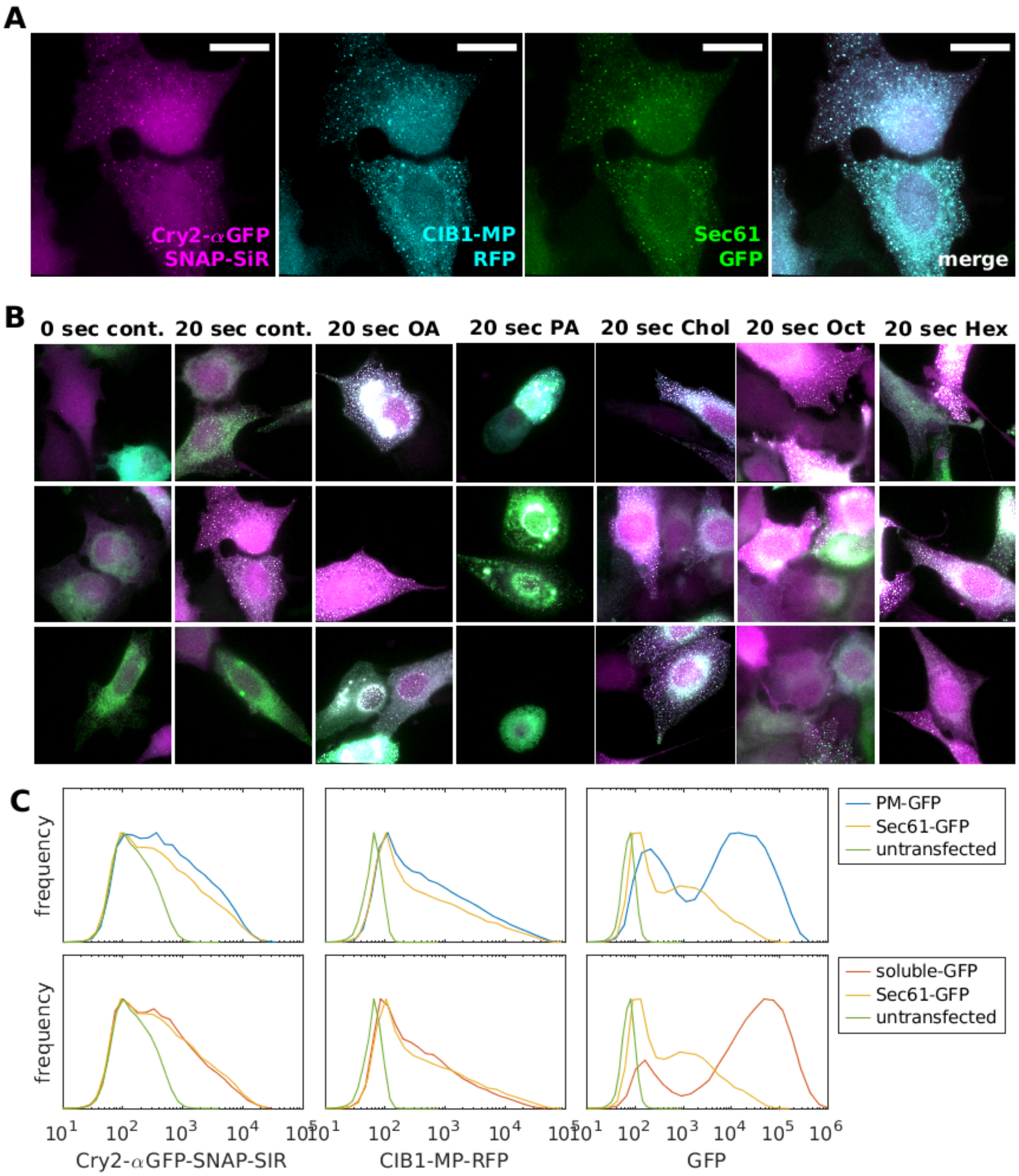


**Supplementary Figure S15: Characterization of U2OS cells expressing Cry2-αGFP and CIB1-MP and Sec61 GFP.** A) Representative epifluorescence image of U2OS cell exposed to 20s of blue light prior to fixation and imaged in three color channels. B) Merged images of representative cells used to quantify cells with clusters under a range of conditions described in the main text. C) Expression level profiles of cells expressing Cry2-αGFP-SNAP labeled with a SiR-SNAP ligand, CIB1-MP-RFP, and one of either PM-GFP, soluble-GFP or Sec61-GFP.

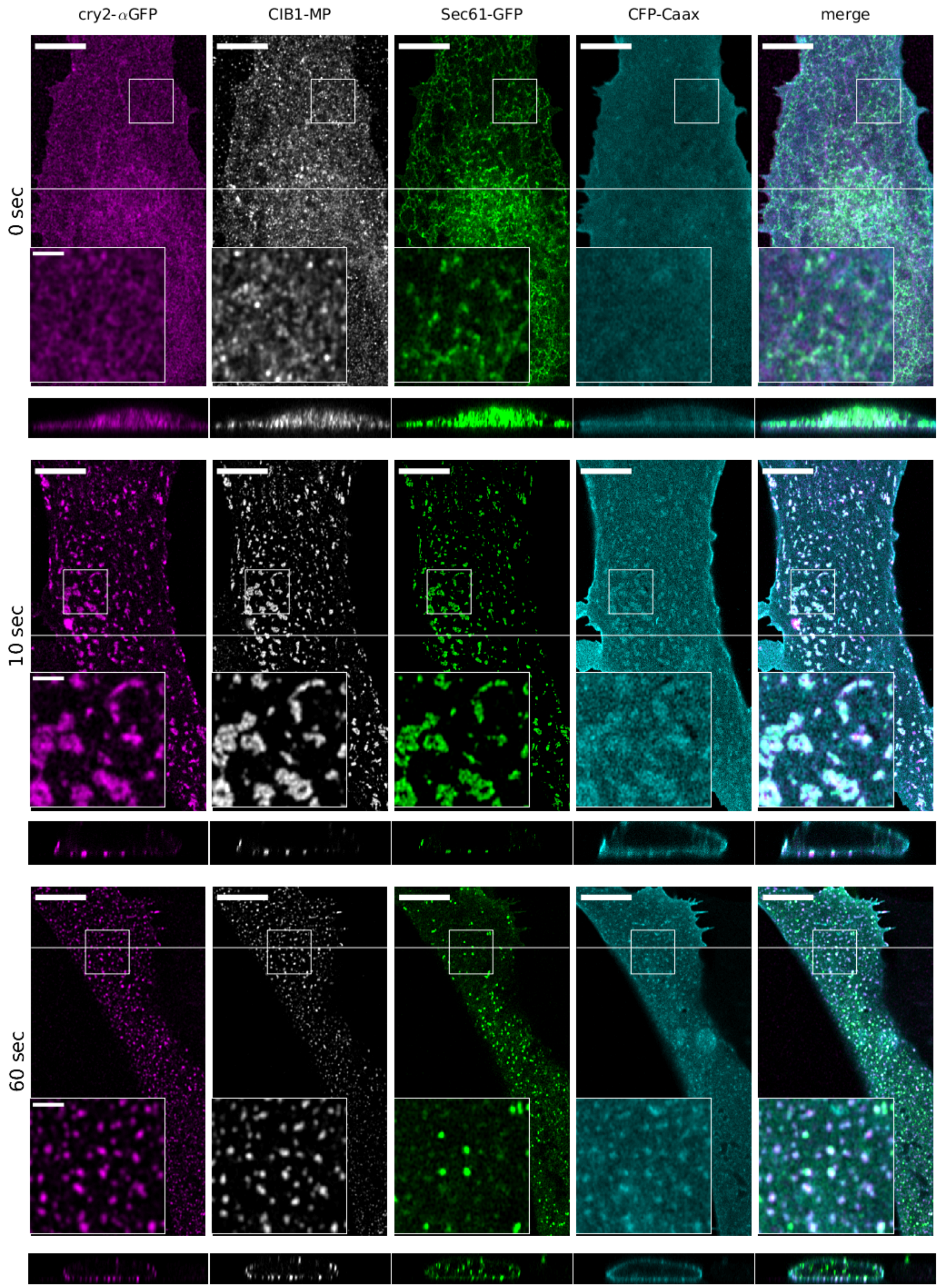


**Supplementary Figure S16: Localization of expressed proteins prior to blue light exposure obtained by Airyscan confocal imaging.** The merged image does not include the CIB1-MP channel.

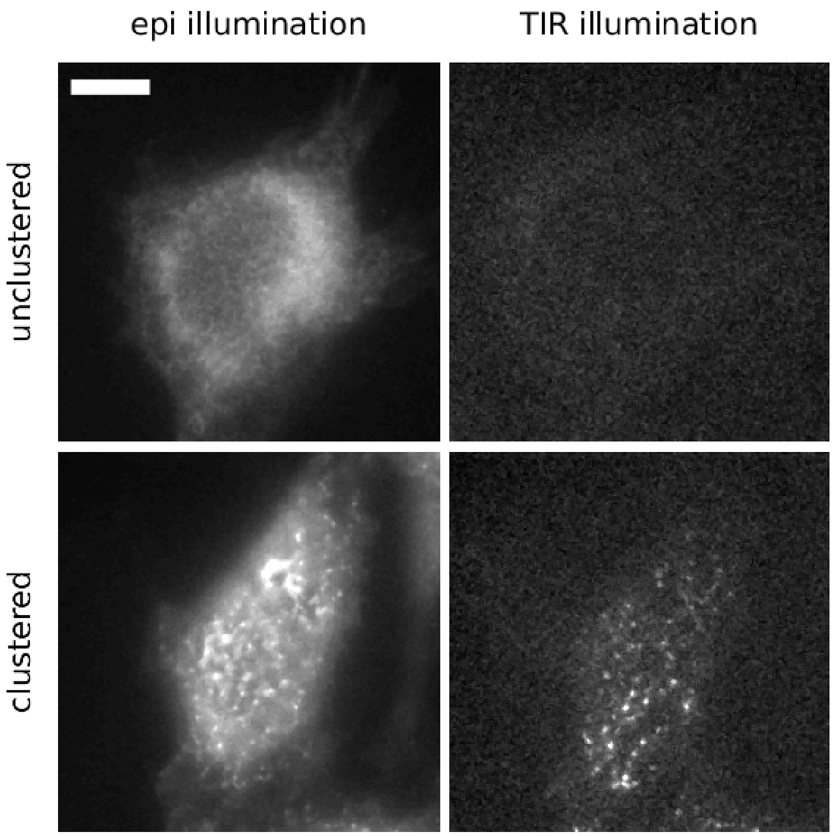


**Supplementary Figure S17: Sec31-GFP puncta are visualized with total internal reflection (TIR) illumination.** U2OS cells expressing Cry2-αGFP,CIB1-MP, Sec61-GFP and CFP-Caax chemically fixed after 5s of illumination with blue light. Dishes contain a mixture of cells that are unclustered and clustered under this condition. (Top) Sec61-GFP imaged under (left) epifluorescence (epi) illumination and (right) TIR illumination in an unclustered cell. No meaningful intensity of the ER membrane label is detected with TIR. (Bottom) Sec61-GFP imaged under (left) epifluorescence (epi) illumination and (right) TIR illumination in a cell exhibiting clusters. Puncta are visualized under TIR illumination, indicating that the ER marker is within several hundred nm of the glass surface.

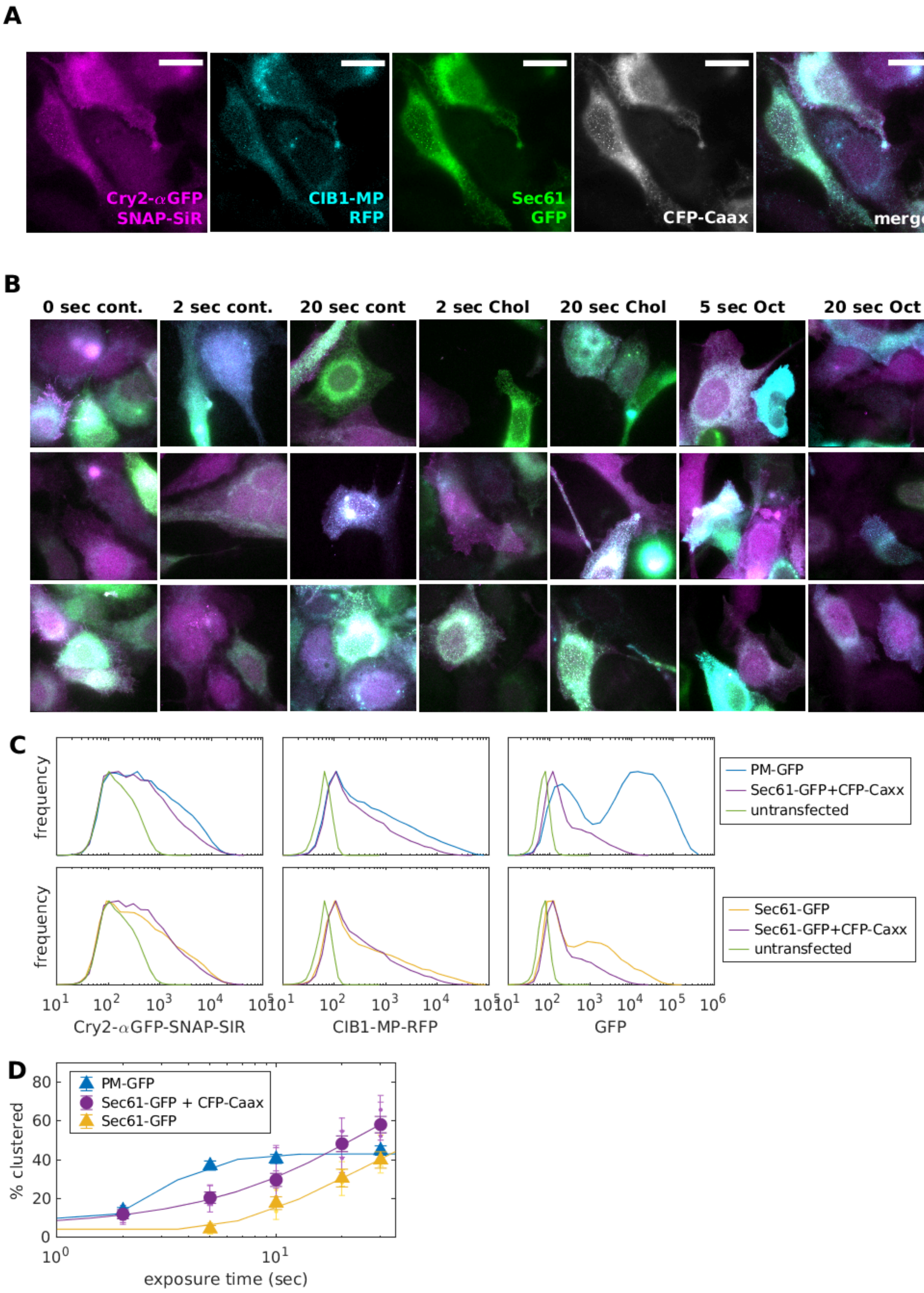


**Supplementary Figure S18: Characterization of U2OS cells expressing Cry2-αGFP and CIB1-MP, Sec61-GFP and CFP-Caax.** A) Representative epifluorescence image of U2OS cell exposed to 20s of blue light prior to fixation and imaged in four color channels. The merged image excludes CFP-Caax. B) Merged images of representative cells, again excluding CFP-Caax, used to quantify cells with clusters under a range of conditions described in the main text. C) Expression level profiles of cells expressing Cry2-αGFP-SNAP labeled with a SiR-SNAP ligand, CIB1-MP-RFP, and either PM-GFP, Sec61-GFP, or Sec61-GFP and CFP-Caax. D) Curves representing the percentages of cells exhibiting puncta as a function of blue light exposure. Cells express Cry2-αGFP, CIB1-MP in addition to the proteins indicated in the legend. Note that expression levels of proteins vary between conditions.

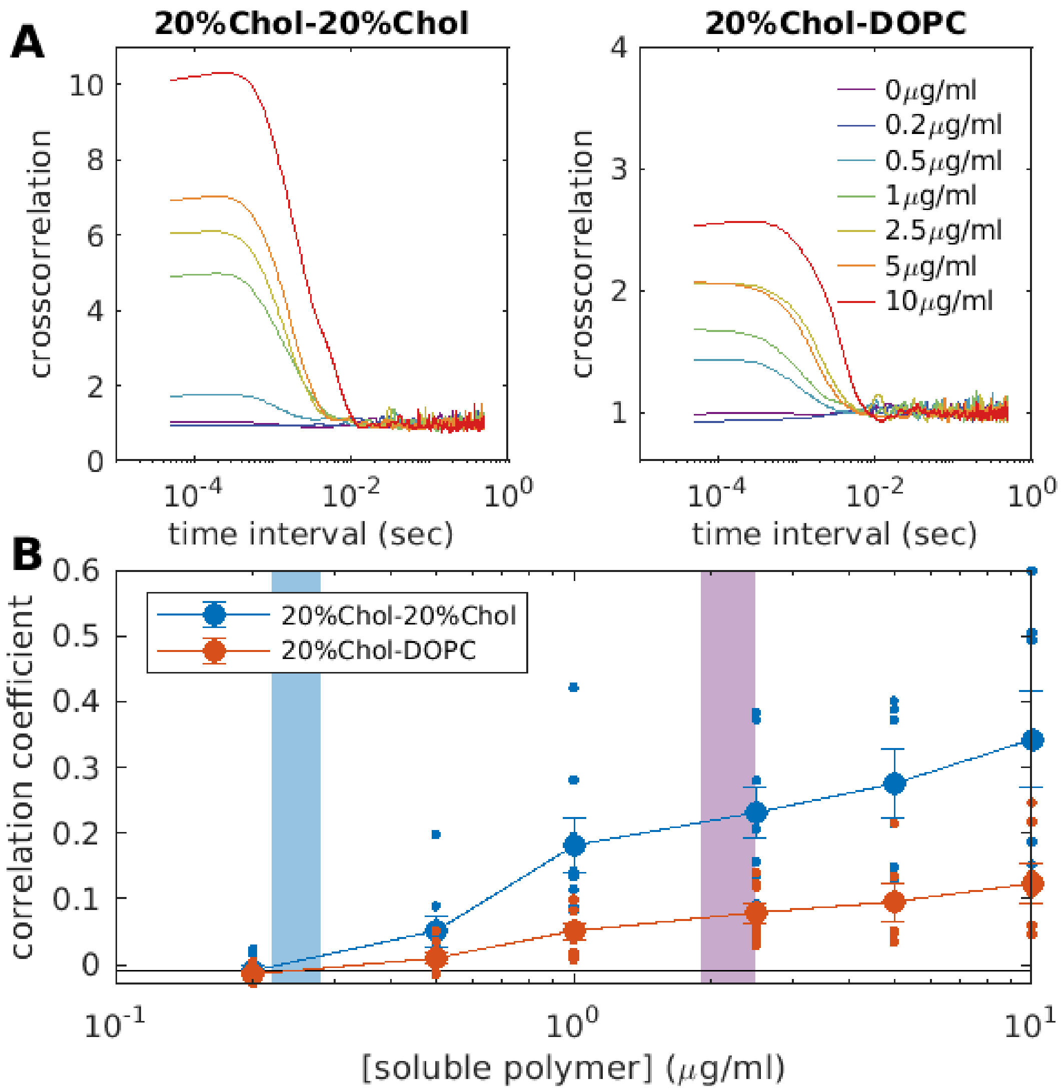


**Supplementary Figure S19: Vesicle adhesion can occur below $c_{prewet}$ in mixed systems.** LUVs containing DOPC, DPPC, 20% Chol, and 4% DOPE-pLys were either labeled with DiO or DiI, and DOPE+4% DOPE-pLys were labeled with DiD, then all three populations were mixed in a sample chamber at 37°C and interrogated by FCCS. Separate measurements were conducted to probe the codiffusion of vesicles labeled containing DiO with DiI (20%Chol-20%Chol) and DiO with DiD (20%Chol-DOPC) at each soluble polymer concentration. A) Crosscorrelation curves acquired from a representative experiment for the titration of soluble pLys/pGlu polymers indicated in the legend. B) A summary of cross-correlation coefficients acquired from 8 separate experiments spanning 4 vesicle preparations. Values obtained from single experiments are represented as small circles, and average values with SEM error bounds are presented as large symbols. The values of $c_{prewet}$ for 20%Chol and DOPC membranes are represented by blue and red shaded regions respectively. These curves demonstrate that DOPC vesicles co-diffuse with 20%Chol vesicles below $c_{prewet}$ of DOPC membranes in a mixed system.

**Supplementary Table 1: Parameters for Simulations shown in Figures**

| | $J_{bulk}$ ($k_BT$) | Membrane Composition | Tether Concentration | $\mu_{bulk}$ , $k_BT$ ($\phi_{bulk}$) | $T_{mem}$ / $T_{c,mem}$ |
|---|---|---|---|---|---|
| Figure 1I | 1.6 | 1 | 0.04 | -4.0, -3.1, -2.9, (0.05) | N/A |
| Figure 1J | 1.6 | 1 | 0.04 | -3.1, -2.9 | N/A |
| Figure 2C | 1.6 | 1, 0.5, 0.5 | 0.04 | -2.9 | N/A, 1.25,0.9 |
| Figure S2A,B | 1.6 | 1, 0.5, 0.5 | 0.04 | -2.9 | N/A, 1.25, 0.9 |
| Figure S2C | 1.6 | 1, 0.5 | 0.04 | [-6,-2.5) | N/A, 1.05 |
| Figure S4 | 1.55 | 1, 0.5 | 0.04 | [-6, -2.5) | N/A, 1.05 |

**Supplementary Table 2: Number of independent replicates shown in Figures**

| **Dataset(s)** | **Figure(s) where it appears** | **Number of independent replicates** |
|---|---|---|
| Cry2-αGFP/CIB1-MP/PM-GFP 22°C | 3D,E,F | 7 |
| Cry2-αGFP/CIB1-MP/GFP 22°C | 3D,E & 4D | 3 |
| Cry2-αGFP/CIB1-MP/PM-GFP 4°C, 37°C | 3D | 3, 3 |
| Cry2-αGFP/CIB1-MP/GFP 4°C, 37°C | 3D | 2, 2 |
| Cry2-αGFP/CIB1-MP/PM-GFP + oct, hex, DMSO | 3D | 3, 3, 7 |
| Cry2-αGFP/CIB1-MP/GFP + oct, hex, DMSO | 3D | 3, 3, 3 |
| Cry2-αGFP/CIB1-MP/PM-GFP + M□CD, M□CD-Chol, A23187 | 3E | 3, 3, 3 |
| Cry2-αGFP/CIB1-MP/Sec61-GFP | 4D | 2 |
| Cry2-αGFP/CIB1-MP/Sec61-GFP + oct, hex, DMSO, M□CD-Chol | 4E | 3, 3, 3, 3 |
| Cry2-αGFP/CIB1-MP/Sec61-GFP | 4E | 3 |
| Cry2-αGFP/CIB1-MP/Sec61-GFP + BSA 3h, 24h | 4F | 3, 3 |
| Cry2-αGFP/CIB1-MP/Sec61-GFP + OA 3h, 24h | 4F | 3, 3 |
| Cry2-αGFP/CIB1-MP/Sec61-GFP + PA 3h, 24h | 4F | 3, 3 |
| DOPC + 4% DOPE-pLys DLS, FCCS | 5C | 3, 2 |
| DOPC + 4% DOPE-pLys DLS, FCCS | 5C | 3, 2 |
| DOPC/DPPC+20% Chol + 4% DOPE-pLys DLS, FCCS | 5C | 3, 2 |
| DOPC+4% DOPE-DBCO DLS, FCCS | 5C | 3, 2 |
| Cry2-αGFP/CIB1-MP/Sec61-GFP/CFP-Caax | 5F | 3 |
| Cry2-αGFP/CIB1-MP/Sec61-GFP/CFP-Caax + oct, DMSO, M□CD-Chol | 5F | 4, 4, 3 |
